# LongPhase-S: purity estimation and variant recalibration with somatic haplotying for long-read sequencing

**DOI:** 10.1101/2025.11.20.689492

**Authors:** Ming-En Ho, Zhenxian Zheng, Ruibang Luo, Huai-Hsiang Chiang, Yao-Ting Huang

## Abstract

Accurate detection of somatic variants is crucial for precision oncology, and long-read sequencing offers unprecedented advantages in resolving complex cancer genomes. However, most long-read somatic callers rely on phasing built for a diploid genome, an assumption violated by various contamination, subclonal heterogeneity, and aneuploidy in tumors. We present LongPhase-S, a novel method that jointly reconstructs somatic haplotypes, infers tumor purity, and recalibrates somatic variants in a purity-aware manner for paired tumor-normal long-read sequencing. By anchoring each somatic read to a parental germline lineage, LongPhase-S provides a phase-resolved view in which germline and somatic reads are disentangled across the genome. Building on somatic haplotyping, LongPhase-S trains a phase-aware purity estimator that outperformed existing methods. Using eight benchmark datasets comprising six cancer cell lines, including breast, melanoma, and lung cancers, LongPhase-S boosted the accuracy of state-of-the-art somatic callers wuth the estimated purity and somatic haplotypes. Specifically, mean F1 scores increased by 4.5% and 7.1% for single-nucleotide variants and insertions and deletions with ClairS, and by 1.2% and 0.5% with DeepSomatic. Collectively, these results showed that somatic haplotyping is a critical yet missing piece in existing somatic callers, which enables purity-aware and phase-resolved variant interpretation in heterogneous tumors.

## 1 Introduction

Cancer arises through the progressive accumulation of somatic genetic alterations that subvert normal cellular programs and enable malignant transformation. Systematic cataloging of these alterations by next-generation sequencing (NGS) has reshaped our understanding of tumor biology and underpins precision oncology. Accurate detection of somatic variants is clinically consequential because specific mutations function as diagnostic and prognostic biomarkers and as actionable therapeutic targets. To maximize specificity and interpretability, most studies adopt a paired tumor-normal design in which DNA from the tumor and a matched normal sample (e.g., blood) are sequenced. The comparison between tumor and normal samples removes inherited, sample-specific germline variants, yielding a high-confidence set of tumor-specific mutations [1, 2]. Nevertheless, because cancer genomes are enriched for complex rearrangements,aneuploidy, and subclonality, the accurate identification of somatic variants remains challenging [3, 4].

Short-read sequencing has predominated in somatic variant discovery owing to its high throughput and cost-effectiveness. However, it is unable to reliably identify mutations within long, repetitive, and structurally complex regions. In recent years, long-read sequencing overcomes these challenges by generating reads that are tens of kilobases in length, enabling improved alignment, haplotype phasing, and direct resolution of complex rearrangements [5, 6]. Building on these advances, deep neural networks trained on long-read sequencing have demonstrated competitive or superior accuracy for calling somatic single-nucleotide variants (SNVs) and insertions/deletions (indels), particularly in difficult regions. For instance, ClairS, created by the developers of Clair3, can identify somatic variants with high accuracy from long reads using pileup- and alignment-based neural networks [7, 8]. Google’s DeepSomatic extends the DeepVariant framework to tumor-normal long-read sequencing by training a convolutional neural network (CNN) using multiple real cancer cell lines [9–11].

Long-read sequencing offers a key advantage for somatic variant detection by enabling robust read-based haplotype phasing. True emerging mutations tend to reside on the same parental haplotype, whereas spurious sequencing errors are haplotype-incoherent and randomly distribute across both [12]. Therefore, phasing long reads prior to somatic variant calling significantly improves the variant detection accuracy. For instance, ClairS employs phased reads by LongPhase and integrates into its full-alignment neural network during somatic variant calling [8, 13]. DeepSomatic implements a tailored phasing strategy within its pipeline and encodes the haplotypes into its CNN [9, 10]. Severus leverages WhatsHap and LongPhase to phase long reads prior to detecting somatic structural variations [14]. Collectively, these independent strategies consistently demonstrate that long-read phasing substantially improves the reliability of somatic variant calling.

However, current phasing approaches assume a single diploid genome, whereas tumor cells comprise mixtures of subclones with variable purity and aneuploidy [15]. In particular, low variant-allele frequencies driven by intra-/inter-tumor heterogeneity, together with sequencing artifacts and limited coverage, reduce the accuracy of phasing [16, 17]. Moreoever, tumor-in-normal contamination further dilute or replicate somatic alleles across paired samples, yielding ambiguous phasing results and inflating both false negatives and false positives [18]. Consequently, clone-specific haplotypes can be collapsed, reads misassigned, and subclonal variants may be mispredicted as sequencing errors. Together, these constraints limit the sensitivity and specificity of existing phasing methods and motivate approaches that can simultaneously estimate tumor purity, normal contamination, and tumor subclonality using long-read haplotype data from both tumor and normal samples.

Here we present LongPhase-S, a novel method for joint reconstruction of somatic haplotypes, estimation of tumor purity, and somatic variant recalibration for paired tumor-normal long-read sequencing. By anchoring somatic reads to their parental germline haplotypes, LongPhase-S disentangles germline and somatic haplotypes in tumor samples and enables a novel haplotype-based estimator for tumor purity. Using the estimated purity and somatic haplotagging reads, LongPhase-S improves the F1-score of somatic SNVs and indels of ClairS and DeepSomatic by on average 4.5% and 1.2% for SNVs and 7.1% and 0.5% for indels, respectively.

## 2 Results

### 2.1 LongPhase-S: purity estimation and variant recalibration with somatic haplotying

LongPhase-S leverages long-read sequencing of a tumor and matched normal to reconstruct somatic haplotypes, estimate purity directly from haplotypes, and perform purity-aware somatic variant recalibration. The components and workflow of LongPhase-S are shown in Figure 1 (see Methods). Briefly, reads are first phased and haplotagged using germline and somatic variants to form germline (HP1/HP2) and somatic haplotypes (HP1-1/HP2-1)(Figure 1(A)). The somatic haplotagging reads carry the somatic mutations and preserve the clonal relationship between the germline and somatic lineages, where HP1-1 and HP2-1 represent the somatically altered descendants of HP1 and HP2, respectively.

**Fig. 1:**
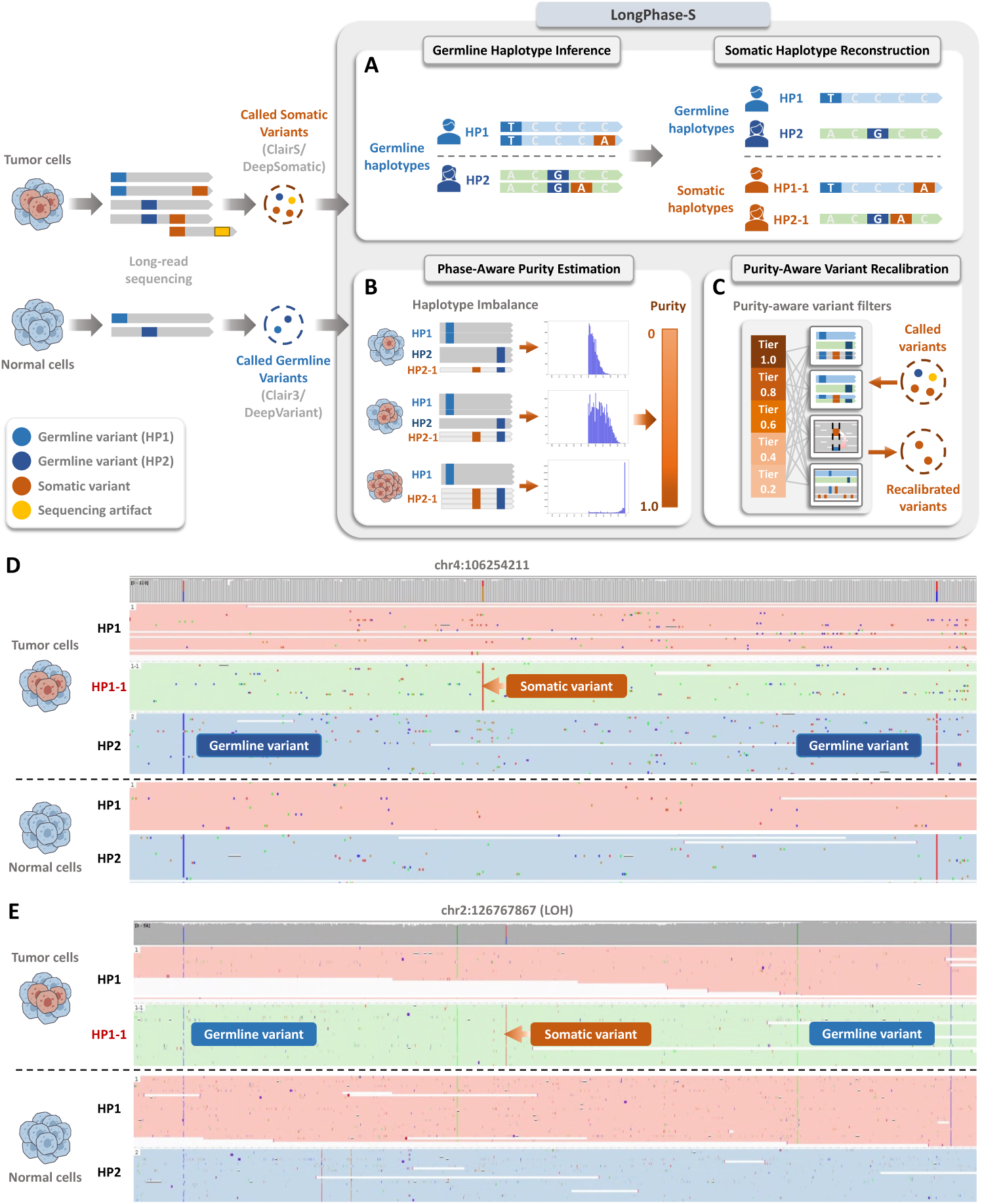
Overview of the LongPhase-S pipeline. (A) Tumor and matched-normal long reads are phased into germline (HP1/HP2) and their descendant somatic haplotypes (HP1-1/HP2-1). (B) The haplotype imbalance between HP1 and HP2 reads correlates with tumor purity, where greater skew indicates higher purity. (C) The inferred purity and somatic haplotagging reads can recalibrate somatic variants by existing callers. (D) The somatic (HP1-1) and germline (HP1/HP2) reads in a region containing a known (SEQC2) somatic vaiant are disentangled in Integrated Genome Viewer (IGV). (E) The somatic reads are correctly tagged as HP1-1 within a large LOH region of HP2-deleted haplotype.

These somatic haplotagged reads provide an indirect yet robust estimate of tumor purity by quantifying haplotype imbalance: the shift in haplotype abundance between HP1 and HP2 reads (Figure 1(B)). In low-purity tumor samples, where reads are dominated by normal cells, the HP1- and HP2-tagged reads are nearly balanced across the genome. As tumor purity increases, the somatic reads (HP1-1 or HP2-1) become over-represented, disrupting the balance of two germline haplotypes by an amount proportional to the tumor cell fraction. By aggregating this haplotype imbalance across the genome with a statistical model, LongPhase-S yields a robust and direct purity estimate without requiring external copy-number profiles.

Using the estimated purity and somatic haplotagging reads, LongPhase-S further recalibrates somatic variant calls of existing callers (e.g., ClairS and DeepSomatic) (Figure 1(C)). Variants that co-segregate with a somatic haplotype and match the expected purity are retained, whereas candidates that disperse across haplotypes or deviate from purity expectations are discarded. This purity-aware variant recalibration improves the F1-score of both somatic SNVs and indels of ClairS and DeepSomatic.

Collectively, LongPhase-S delivers a read-level, phase-resolved view in which germline and somatic reads are disentangled across long tracts of the genome. In a paired tumor-normal sequecing dataset (HCC1395_HKU), reads harboring the somatic variant (i.e., HP1-1) were accurately anchored to their ancestral haplotype and separated from the remaining germline reads (Figure 1(D)). Within a region of loss of heterozygosity (LOH) where one haplotype (HP2) was deleted, LongPhase-S correctly tagged the somatic reads as HP1-1 instead of either germline haplotype (i.e., HP1 and HP2) (Figure 1(E)). Note that conventional germline phasing tools would misassign somatic reads either to the remaining haplotype (HP1) or to the deleted haplotype (HP2) in LOH regions, because they assume a diploid genome without modeling clonal haplotypes. Therefore, somatic haplotyping with LongPhase-S can provide higher resolution of clonal architecture in a heterogenous tumor sample.

### 2.2 Accurate and robust tumor purity estimation with LongPhase-S

Building on somatic haplotyping, we define the germline haplotype imbalance ratio (GHIR), which quantifies the genome-wide skew of reads toward one germline haplotype (0.5 denotes parity and 1.0 indicates dominance of one haplotype) (see Methods). In two cancer cell lines (HCC1395 and COLO829), the GHIR distribution at 1.0 purity was narrowly concentrated at high values, suggesting a strong genome-wide haplotype imbalance (Figure 2(A)). The progressive addition of normal cells (i.e., reducing purity from 0.8 to 0.2) shifted the median of GHIR toward 0.5 and steadily reduced the dispersion. The same median and dispersion changes of GHIR with tumor purity were consistently observed in breast (HCC1937, HCC1954), melanoma (COLO829), and lung (H1437, H2009) cancer cell lines (Supplementary Figure 1, Supplementary Table 1), suggesting that GHIR captures a conserved relationship between tumor purity and haplotype imbalance across distinct tumor types.

**Fig. 2:**
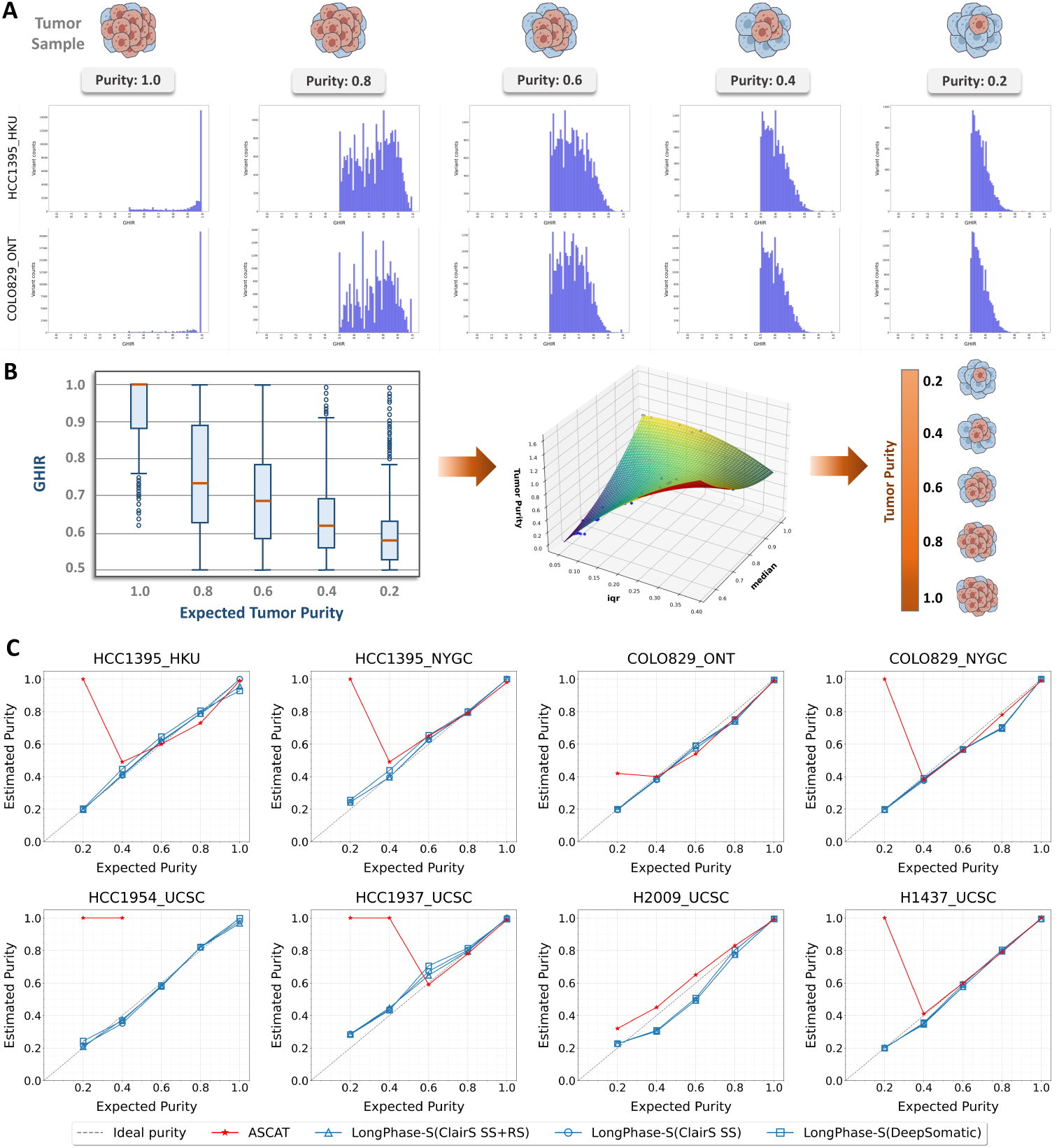
(A) GHIR distributions for the HCC1395_HKU and COLO829_ONT cell lines at tumor purities of 1.0, 0.8, 0.6, 0.4, and 0.2 (left to right). (B) Boxplot summaries of GHIR (median, IQR) were used as features to fit a model for purity estimation. (C) Comparison of purity estimation for LongPhase-S of three configurations (blue lines) and ASCAT (red line) across eight cell line datasets.

These purity-dependent changes in median and dispersion of GHIR motivated a novel, haplotye-based approach for purity estimation (Figure 2B). We trained a polynomial regression model using two GHIR-derived features, the sample median and interquartile range (IQR), along with synthetic tumor-normal mixtures spanning purities 0-1.0 (see Methods). The resulting phase-aware model predicts tumor purity directly using the somatic haplotagging output of LongPhase-S. Cross validations indicated that the model achieved an average *R*^2^ ≈ 0.96 with minima; training-testing differences in terms of MAE, MSE and RMSE, supporting its stability and mitigating overfitting. (Supplementary Figure 2 and Supplementary Table 2).

We then benchmarked LongPhase-S prediction against a well-known purity estimator (ASCAT) using *in silico* mixtures with expected tumor purities in eight cancer cell line datasets [19]. LongPhase-S predicted purity using somatic variants from ClairS (SS+RS model), ClairS (SS model), and DeepSomatic (Figure 2). Across all datasets, the three LongPhase-S configurations produced well-calibrated predictions that closely matched the expected purities at low-to-high levels. In contrast, ASCAT exhibited substantial variability and frequent overestimation, particularly when purity ≤0.4. For instance, it mispredicted purity as 1.0 when 0.2 was expected in six datasets (e.g., COLO829_NYGC). In HCC_1954, ASCAT failed to generate predictions for purity greater than 0.4. Together, these results indicate that LongPhase-S is a highly accurate, robust, and phase-aware method for tumor purity estimation.

### 2.3 LongPhase-S improves somatic SNV and indel calling through variant recalibration

We next assessed whether the variant recalibration by LongPhase-S enhances two top long-read variant callers, ClairS (SS+RS model) and DeepSomatic. Each caller was benchmarked both in its standalone form and in combination with LongPhase-S, yielding four configurations across eight cancer cell-line datasets: ClairS, ClairS+LongPhase-S, DeepSomatic, and DeepSomatic+LongPhase-S. Applying LongPhase-S to recalibrating ClairS SNVs consistently increased recall in all datasets (mean: +4.5%, +1.4% to 14.4%, e.g., 0.78 to 0.93 in HCC1937_UCSC) (Figure 3(A)). These recall gains were accompanied by modest reductions in precision (mean:-1.3%, -3.9% to +0.8%, e.g., 0.95 to 0.91 in HCC1395_HKU). Consequently, the net effect was an F1-score improvement of ClairS in 6/8 datasets (mean: +1.9%, - 0.3% to +8.3%), with notable gains in HCC1395_HKU (+8.3%) and HCC1954_UCSC (+5%). When applied to DeepSomatic, LongPhase-S primarily enhanced precision (mean: 3.1%, +0.2% to +17.2%, e.g., 0.75 to 0.93 in HCC1395_HKU), while recall slighly decreased (mean: -0.5%, -0.1% to -1%). Consequently, F1-score changes were small (mean: +1.2%, -0.3% to +7.8%), yielding only modest gains in 6/8 datasets and marginal declines in the others.

**Fig. 3:**
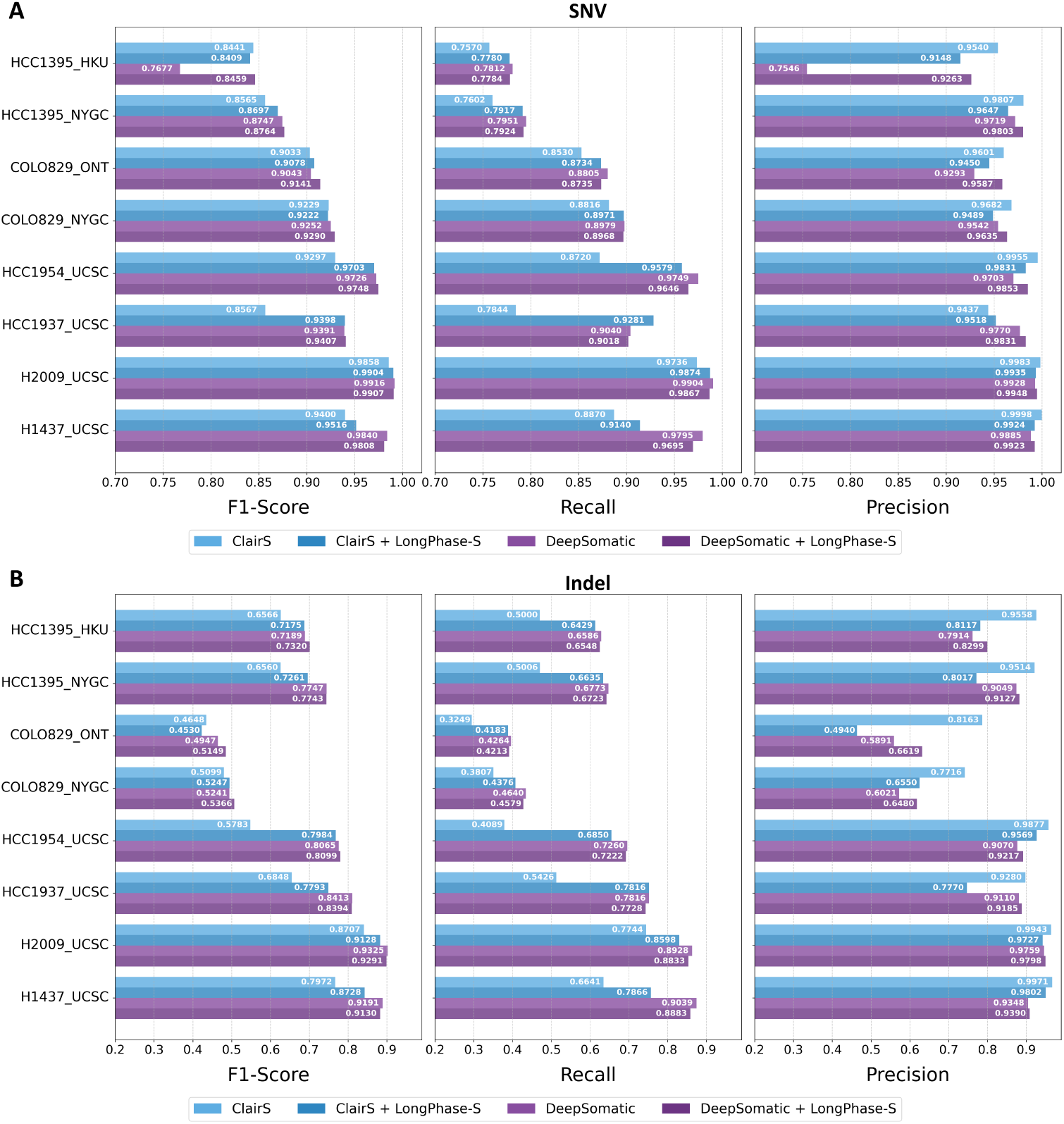
F1, recall, and precision of adding LongPhase-S to recalibrate somatic variants of ClairS and DeepSomatic using eight cancer cell line datasets for (A) SNV and (B) indel calling.

The performance of indel calling by adding LongPhase-S to ClairS and DeepSomatic exhibited a similar trend (Figure 3(B)). When applied to ClairS, LongPhase-S substantially increased recall across all eight datasets (mean: +14.7%, +6% to +28%, e.g., 0.41 to 0.69 in HCC1954_UCSC and 0.50 to 0.64 in HCC1395_HKU), accompanied by a decrease in precision (mean: -11.9%, -2% to -32%; e.g., 0.99 to 0.96 in HCC1954_UCSC and 0.96 to 0.81 in HCC1395_HKU). Overall, ClairS+LongPhase-S improved F1-score in 7/8 datasets (mean: +7.1%), including large gains in HCC1954_UCSC (0.58 to 0.80, +22%) and HCC1937_UCSC (0.68 to 0.78, +9.5%). The only exception was COLO829_ONT where F1 decreased slightly (0.46 to 0.45) due to a marked precision drop (0.82 to 0.49) despite higher recall (0.32 to 0.42). For DeepSomatic, LongPhase-S primarily increased precision (mean: +2.4%; up to +7.3% in COLO829_ONT and +4.6% in COLO829_NYGC) while slightly reducing recall (mean: −0.7%). Consequently, F1 changes for DeepSomatic were modest (mean: +0.5%), with small gains in about half of the datasets (e.g., HCC1395_HKU 0.72 to 0.73; COLO829_ONT 0.49 to 0.51) and marginal declines in the rest (e.g., H1437_UCSC 0.919 to 0.913).

### 2.4 Performance comaprison of somatic SNV and indel calling in varying purity

We further evaluated the performance of adding LongPhase-S to ClairS and DeepSomatic under varying tumor purity (0.2-1.0)(Figure 4(A)). For ClairS, LongPhase-S delivered the largest F1 gains at low-intermediate purity (e.g., 0.45 to 0.72 in COLO829_NYGC at 0.2 purity). For HCC1937_UCSC and HCC1954_UCSC, LongPhase-S significantly improved F1 of ClairS at all purity (e.g., ∼10% gains in all purity levels of HCC1937_UCSC). The improvment of ClairS was mainly driven by substantial increases in recall (e.g., up to 22% in HCC1395_HKU at 0.2 purity) (Supplementary Figures 3). The precision was often not offset by the recall gain and occasionally increased at lower purity (e.g., 6% in HCC1954_UCSC at 0.2 purity) (Supplementary Figure 4). For DeepSomatic, LongPhase-S primarily increased precision at higher purity levels (e.g., 0.83 to 0.95 in HCC1395_HKU at 1.0 purity) (Supplementary Figures 3 and 4). The recall on DeepSomatic were slighly reduced (less than 1% in all datasets) in all datasets. Overall, the F1 gains of adding LongPhase-S to DeepSomatic were marginal compared with those for ClairS (≤l1%). The improved accuracy of ClairS by LongPhase-S was often on par with DeepSomatic alone. Therefore, the improvement conferred by LongPhase-S is consistent among different cell lines and different somatic callers. Gains for ClairS were largely recall-driven at low to medium purity, whereas improvements for DeepSomatic were precision-driven at higher purity.

**Fig. 4:**
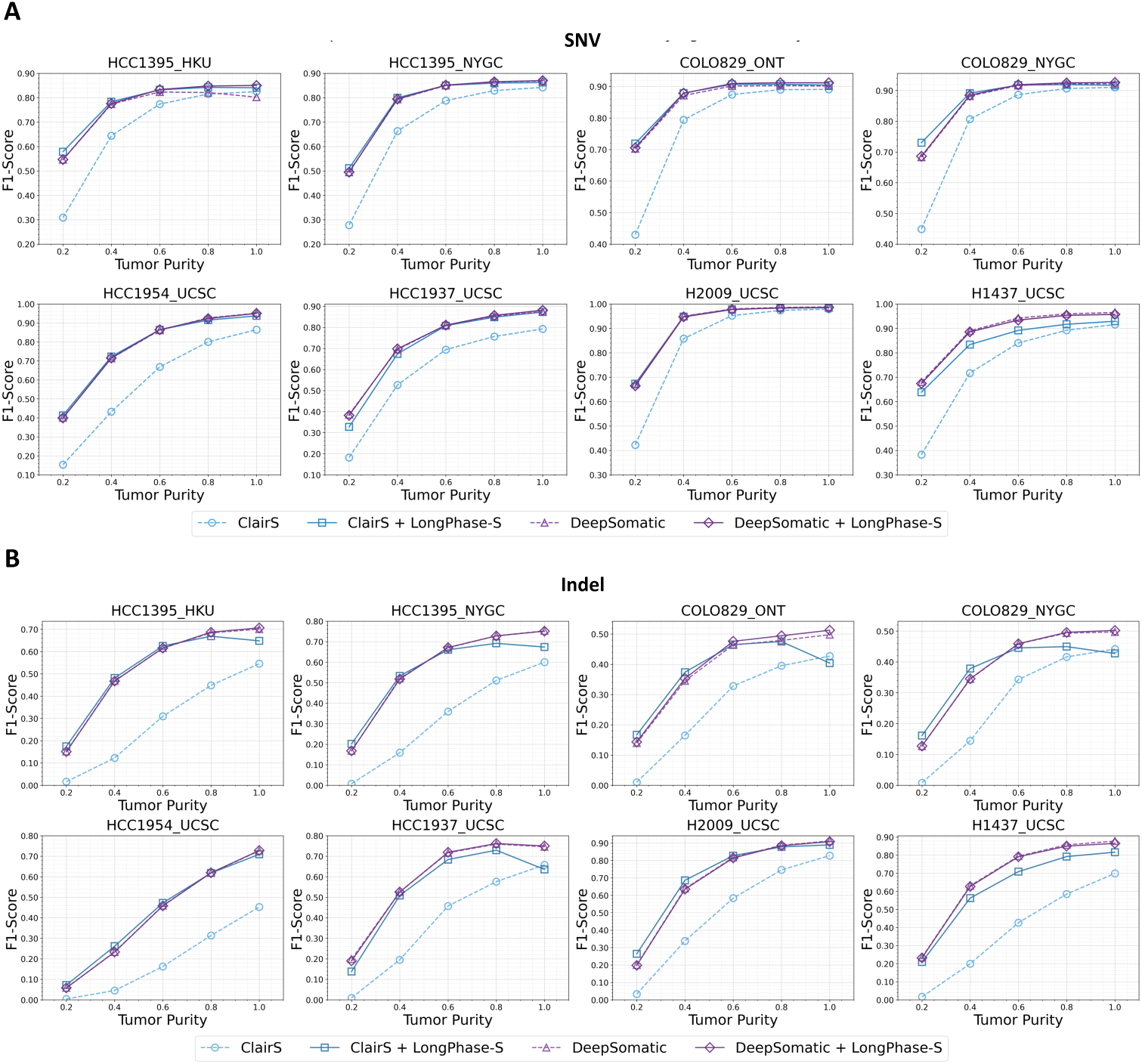
The F1-scores of adding LongPhase-S to recalibrate somatic variants of ClairS and DeepSomatic across varying tumor purity and cell lines for (A) SNV and (B) indel calling.

When recalibrating somatic indels (Figure 4(B)), adding LongPhase-S to ClairS produced large F1 gains mainly at low-intermediate purity across all cell lines, typically by ∼+15-30%. Examples include COLO829_NYGC at 0.2 purity (F1 from 0.01 to 0.15) and H2009_UCSC at 0.4 (F1 from 0.35 to 0.67). These improvements were primarily recall-driven. e.g., HCC1937_UCSC recall increased from ∼0.10 to 0.39 at 0.4 and from 0.52 to 0.61 at 1.0 purity (Supplementary Figure 5). The precision mostly held unchanged at lower purity and decreased at higher purity in certain datasets (e.g., HCC1395_HKU 0.91 to 0.67 and HCC1937_UCSC 0.9 to 0.55 at 1.0)(Supplementary Figure 6). In contrast, adding LongPhase-S to DeepSomatic had only marginal impact on F1 (generally ≤+1-2%). The recall increased modestly at intermediate purity (e.g., H1437_UCSC 0.57 to 0.68 at 0.6) with slight reduction of precision (e.g., H2009_UCSC 0.98 to 0.97 at 0.6-1.0). Overall, for somatic indel recalibration, LongPhase-S substantially boosts ClairS by bringing its accuracy close to DeepSomatic, and its incremental effect on DeepSomatic leaves DeepSomatic’s performance largely unchanged.

To assess whether somatic haplotagging can be exploited within existing callers rather than via post-recalibration, we applied LongPhase-S to anchoring somatic reads using the initial variant candidates generated by two pileup model (SS and SS+RS) of ClairS. The somatic variants with reads tagged by LongPhase-S exhibitied higher accuracy than those produced by ClairS pileup models (e.g., an ∼18% F1 gain in HCC1395_HKU at 1.0 purity for the SS pileup model) (Supplementary Figure 7). The recall of ClairS and LongPhase-S were not much different in most datasets (Supplementary Figure 8), but the precision of ClairS was greatly reduced at higher purity while LongPhase-S was less affected (e.g., ∼50% to ∼80% in HCC1395_HKU at 1.0 purity) (Supplementary Figures 8 and 9). When comparing the two internal models within ClairS, the SS model degraded at higher purity but benefited substantially more by LongPhase-S haplotagging than the SS+RS model. Consequently, the somatic variants with reads haplotagged by LongPhase-S are more accurate than the initial variant candidates, which can be embedded into its subsequent (full-alignment) model. This result establishes that phase information derived from somatic haplotagging is the key differentiator. It is therefore imperative for existing variant callers to move beyond germline haplotypes and integrate somatic haplotype awareness as an essential component of their calling algorithms.

### 2.5 Read-level benchmark of somatic haplotagging accuracy

Because no benchgmark existed for evaluating germline and somatic haplotagging, we first established a read-level truth set of haplotagging using known somatic varaints (see Methods). This benchmark is then used to evaluate the accuracy of somatic reads tagged by LongPhase-S over DeepSomatic and ClairS. For somatic reads anchored with germline haplotype background (HP1-1 and HP2-1), LongPhase-S achieved uniformly high accuracy using DeepSomatic variant calls (Figure 5(A)). The F1-scores were ≥0.98 in six of eight datasets (e.g., F1 = 0.99 in HCC1395_NYGC) with both high recall and precision. In contrast, the tagging accuracy of somatic reads lacking germline haplotype support (HP3) were substantially lower (e.g., F1 = 0.746 for HP3 v.s. 0.929 for HP1-1 in HCC1937_UCSC) due to significantly reduced recall (e.g., recall = 0.596 for HP3 v.s. 0.871 for HP1-1 in HCC1937_UCSC). The tagging precision was consistently high for all three types of somatic tagging reads (e.g., ≥0.98). The accuracy of LongPhase-S somatic haplotagging on ClairS showed slightly lower F1 scores for HP1-1/HP2-1 reads compared with DeepSomatic (e.g., HP1-1 F1 = 0.880 with ClairS vs. 0.926 with DeepSomatic in HCC1954_UCSC) and diminished recall for HP3 (e.g., recall = 0.719 with ClairS in H1437_UCSC). Collectively, these results indicate that LongPhase-S provides higher accuracy for HP1-1/HP2-1 reads than HP3.

**Fig. 5:**
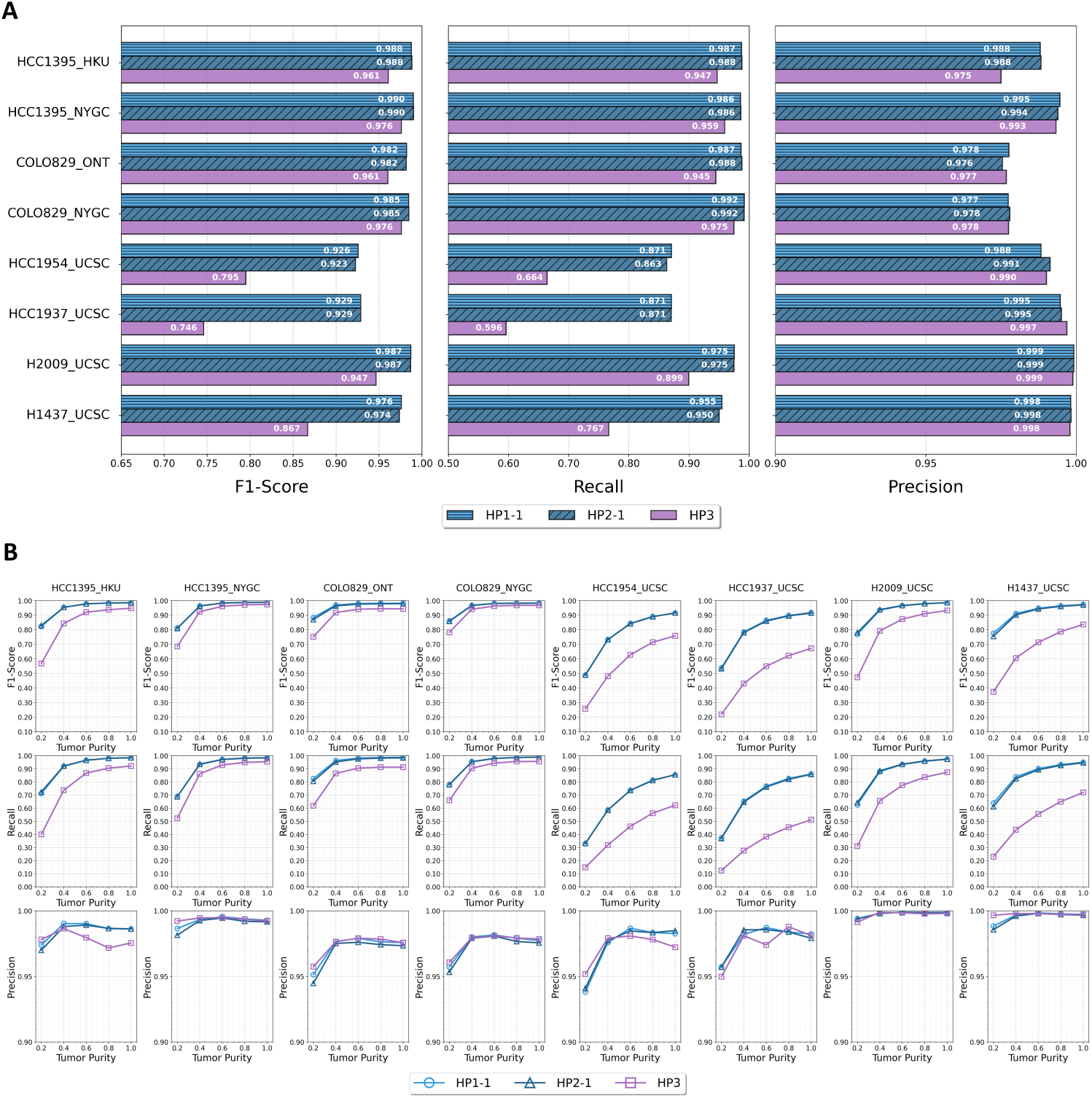
Read-level accuracy of somatic haplotagging (HP1-1, HP2-1, and HP3) by DeepSomatic+LongPhase-S across (a) eight cancer cell line datasets and (B) varying tumor purities.

The accuracy of somatic haplotagging by LongPhase-S was further evaluated under varying tumor purities for DeepSomatic and ClairS (SS and SS+RS models) (Figure 5(B)). The F1-scores of LongPhase-S haplotagging were continuously improved as the tumor purity increased, which were mainly driven by the larger recall while precision was steadily above 0.95 regardless of purity. For instance, at purities of 0.6-1.0, LongPhase-S exhibited high precision and recall for HP1-1 and HP2-1 in HCC1395_HKU (both *>*0.95). However, recall dropped sharply to 0.6 at a 0.2 purity, albeit precision remained high. Similarly, the F1 scores of HP1-1/HP2-1 reads are higher than those of HP3 across all purity levels on all datasets. The haplotagging accuracy of LongPhase-S on ClairS (SS and SS+RS models) exhibited simlartreands (Supplementary Figures 10 and 11). Overall, these results indicate that somatic haplotagging accuracy followed a similar pattern to that of somatic SNV and indel recalibration, where the overall accuracy of HP-1/HP-2 and HP-3 increased with tumor purity and somatic reads with germline support had higher accuracy.

## 3 Discussion and Conclusion

LongPhase-S extends long-read haplotyping into the somatic setting and shows that somatic haplotagging reads can be leveraged to reconstruct somatic haplotypes, infer tumor purity, and recalibrate somatic variant calls. By turning phasing imbalance into a robust purity signal and by enforcing haplotype consistence during variant recalibration, it improves the accuracy of purity estimation and somatic calling. By jointly haplotagging tumor and matched-normal reads into germline and their somatically altered descendants, LongPhase-S provides a read-centric view of heterogenous tumor samples in which clonal relationships are preserved across long genomic regions.

Unlike previous purity estimators that rely on allelic profiles, copy numbers, or gene expression [19–23], our estimation via somatic haplotagging is the first phase-aware method which provides higher accuracy especially at low purity levels. Although our estimation is robust, it can in principle be distorted by large-scale allelic imbalance not attributable to purity, e.g., LOH and aneuploidy. Even many cell lines in our datasets are with chromosome-scale LOH and aneuploidy (e.g., HCC1395), the estimation was still unaffected by these allelic imbalance events. We hypotehsize the robustness of LongPhase-S in the presence of LOH and aneuploidy may stem from phase-aware haplotype estimation, whereas allelic-centric or copy-number methods are more susceptible to these imbalance events.

Purity-aware variant recalibration demonstrates that somatic haplotagging is a critical yet missing piece in existing long-read somatic callers. When applied to ClairS outputs, LongPhase-S consistently increased recall with modest precision trade-offs, yielding net gains in F1 at low-intermediate purity. For DeepSomatic, gains were primarily precision-driven and concentrated at higher purity with little impact on recall. These caller-dependent results may be due to their nerual network differences and their training data. Notably, by applying LongPhase-S to tag the intermediate outputs of ClairS (i.e., the pileup candidates), the F1 improves due to better precision gains with minimal recall loss. This finding suggests that somatic haplotagging reads are likely beneficial to existing callers if directly embedded into the models rather than as a post filter.

Our read-level haplotagging benchmark clarifies when somatic haplotyping is most reliable. The somatic reads which are decendants from a germline haplotype (HP1-1/HP2-1) showed uniformly high precision and recall in most datasets (F1 ≥0.98), and the accruacy improved monotonically with purity. By contrast, somatic reads lacking germline background (HP3) were harder to tag due to lower recall. Without germline anchors, phase must be inferred from sparser somatic linkage, which becomes unreliable at low variant density or in regions depleted of callable germline heterozygotes. Dataset variability further highlights that somatic haplotagging accuracy depends on sample quality, variant density, local mappability, and the complexity of underlying clonal structure.

Our study also has limitations. We trained and evaluated on eight cancer cell line datasets and simulated mixtures, which provide ground truth for purity but may underrepresent the genomic complexity, stromal admixture, and spatial heterogeneity of primary tumors [15, 24, 25]. Subclonal CNAs, LOH, and aneuploidy can create chromosome- or clone-specific allelic skews that partially mimic purity effects [20, 26, 27], although LongPhase-S leverages somatic haplotags to mitigate this, a fully joint model of purity, ploidy, and subclonality would further reduce ambiguity. The integration of orthogonal signals such as methylation phasing may enhance the tagging accuracy of somatic reads lacking germline haplotype support [28]. In summary, LongPhase-S reframes long-read tumor analysis around somatic haplotypes. We anticipate that incorporating somatic haplotypes as features during both inference and learning will enable more accurate, interpretable, and clone-aware cancer genome analyses with long reads.

## 4 Methods

### 4.1 Overview of LongPhase-S

The LongPhase-S workflow comprises four stages: preliminary somatic haplotagging, tumor purity estimation, somatic variant recalibration, and somatic haplotype reconstruction (Figure 6). In the first stage, somatic and germline variants are first called (by ClairS/DeepSomatic and Clair3/DeepVariant) and used to reconstruct the two parental germline haplotypes. Preliminary somatic haplotagging assigns sequencing reads to either germline or somatic haplotypes by leveraging phased germline haplotypes with somatic variants. Second, genome-wide germline haplotype imbalance profiles are computed, low-confidence variants are removed using imbalance- and coverage-based filters, and a polynomial regression model was trained from the cleaned imbalance distributions to estimate tumor purity. Third, this purity estimate guides a tiered set of filters for purity-aware variant recalibration, in which filtering parameters are tuned by a global/local hybrid search to yield a high-confidence set of recalibrated somatic variants. Finally, these recalibrated somatic variants are re-phased to distinguish germline from somaticreads with a tagging inheritance mechanism, providing a phase-resolved view of tumor sequencing data.

**Fig. 6:**
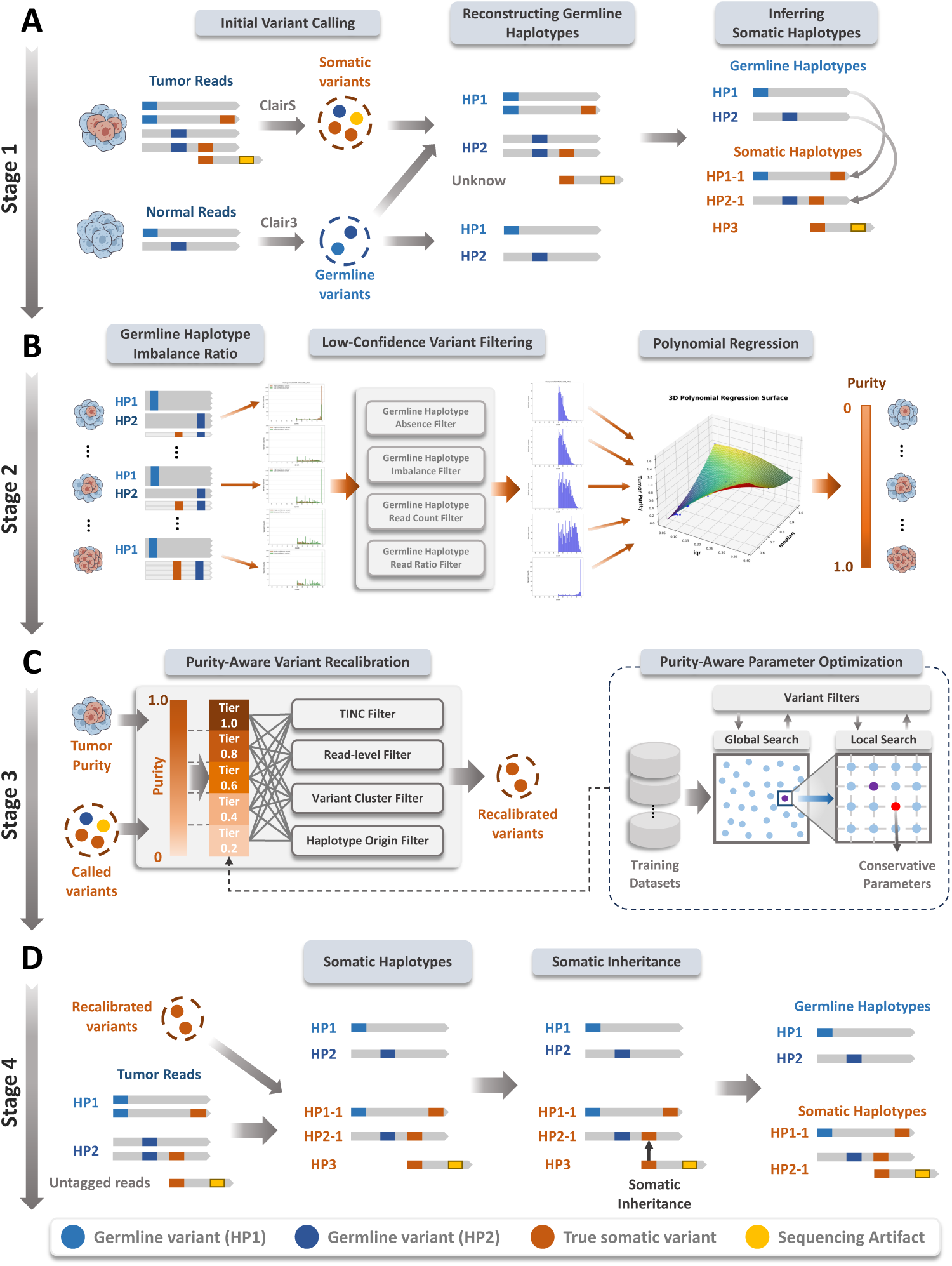
(A) LongPhase-S takes long-read sequencing of tumor and normal samples as input and first calls somatic and germline variants, reconstructs diploid germline haplotypes, and infers initial somatic haplotypes (Stage 1). (B) Genome-wide germline haplotype imbalance profiles are then used to train a polynomial regression model that estimates tumor purity (Stage 2). (C) This purity estimate guides a tiered set of haplotype- and read-level filters whose parameters are tuned on training data to recalibrate somatic variant calls (Stage 3_2_)._3_ (D) Recalibrated variants are phased on tumor reads to assemble somatic haplotypes with a tagging inheritance mechanism (Stage 4).

### 4.2 Preliminary somatic haplotagging

Long-read tumor-normal pairs were aligned to the GRCh38 reference using minimap2 (v2.24-r1122) [29]. Germline variants were called from the normal sample with Clair3 (v1.0.10) or DeepVariant (v1.8.0) [7, 9, 10], and somatic variants were called from the tumor sample with ClairS (v0.4.1) or DeepSomatic (v1.8.0) [8, 11]. We phased the normal-sample germline variants using LongPhase (v1.7.3) [13] to reconstruct the two parental haplotypes (HP1 and HP2).

Tumor reads were then assigned to haplotypes by allelic concordance with the phased germline heterozygotes. Reads carrying only germline variants were tagged as HP1 or HP2. Reads that carried both germline and somatic variants and could be unambiguously traced to HP1 or HP2 were classified as the corresponding somatic subhaplotypes (HP1-1 or HP2-1). When haplotype origin could not be resolved (e.g., in the absence of informative germline variants), reads harboring somatic variants were provisionally assigned to HP3. This procedure provides an initial reconstruction of germline and somatic haplotypes in the tumor sample and establishes a framework for downstream refinement.

### 4.3 Tumor purity estimation

Accurate quantification of the fraction of cancer cells in a sample is critical for sensitive detection of somatic variants and for haplotype phasing, particularly when variant allele frequencies (VAFs) are low. We infer tumor purity directly from preliminary somatic haplotagging by measuring somatic-induced haplotype imbalance in the tumor sample termed Germline Haplotype Imbalance Ratio (GHIR). For each potential somatic variant, GHIR is defined as the ratio between the higher of the two haplotype-specific read counts (HP1 or HP2) and the total read count covering that site. Formally, for variant *i* in the tumor sample,

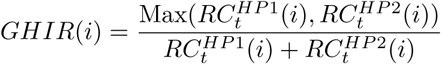

where 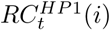 and 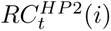 denote the haplotype-specific read counts supporting HP1 and HP2, respectively,, at *V_t_*(*i*) in sample *s*, *t* denotes the tumor sample, and *n* denotes the normal sample.

In normal diploid cells outside regions affected by copy-number variation, parental haplotypes are present in similar proportions, and reads covering heterozygous sites are therefore assigned to HP1 and HP2 with frequencies near 0.5. The GHIR distribution of a normal sample is sharply peaked at 0.5 (Supplementary Figure 12). During tumorigenesis, however, large-scale copy-number alterations, loss of heterozygosity, and aneuploidy frequently disrupt this balance, amplifying or deleting one parental haplotype. Consequently, the GHIR distributions are skewed toward 1 in tumors, with the degree of skew correlating predictably with tumor purity. We exploit the relationship between GHIR and tumor purity by training a statistical model on the GHIR distribution to estimate purity.

In practice, the GHIR distribution is often confounded by a substantial number of false somatic variants called by long-read somatic variant callers (e.g., ClairS and DeepSomatic). Through benchmarking against well-characterized cell lines (e.g., SEQC2 for HCC1395), we found that the variants located in low-confidence regions (defined by SEQC2) disrupt the GHIR distribution. Unfortunately, the SEQC2 high-confidence regions are tailored for a particular cancer cell line (HCC1395), which cannot be used to filter variants of other cancer cell lines. Therefore, we developed a Low-Confidence Variant Filtering (LCVF) module within LongPhase-S for removing low-confidence variants prior to the reconstruction of GHIR and the training of purity-estimating model. Details of LCVF and the model are described below.

#### 4.3.1 Low-confidence variant filtering

Using the matched normal sample as a reference, LCVF first removes somatic loci with insufficient haplotagging reads, discard variants that exhibit pre-existing haplotype imbalance, filters variants under copy-number variations, and ignore variants with insufficient read supports (Supplementary Figure 13). A locus is retained only if it passes all four filters.

##### Minimum Haplotagging Filter

Discard somatic loci with zero haplotype-tagged reads (HP1 or HP2) in the normal or in the tumor samples. Retain only loci with ≥ 1 haplotype-tagged read (HP1/HP2) in both normal and tumor.

##### Germline Haplotype Imbalance Filter

This filter examines the GHIR distribution within the normal sample to remove variants that exhibit a pre-existing haplotype imbalance (Supplementary Figure 14). In a normal sample, a GHIR deviation from 0.5 in a given region suggests the presence of germline structural variations instead of tumor-induced imbalances. Such pre-existing imbalances would also exhibit GHIR skewing. Therefore, a variant is retained only if its GHIR in the normal sample, *GHIR_n_*(*i*), is less than a predefined threshold, 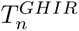, thereby ensuring that the subsequent GHIR distribution focuses on genomic regions truly balanced in the normal genome.

##### Germline Haplotype Read Ratio Filter

The GHIR statistics are vulnerable to incorrect alignment of reads, which is indicative of low-confidence somatic variants. We examine the proportion of reads supporting germline haplotypes at each somatic variant locus using the normal sample as a reference (Supplementary Figure 15). For a given somatic variant locus *V_t_*(*i*), the Germline Haplotype Read Ratio (GHRR) is calculated as:

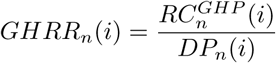

where 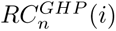 denotes the number of reads assigned to either germline haplotype (i.e., 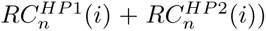, and *DP_n_*(*i*) denotes the total read depth, all measured at *V_t_*(*i*) in the normal sample. By using the SEQC2 benchmark, we found a high proportion of reads support either the HP1 or HP2 germline haplotypes in high-confidence regions, typically resulting in a *GHRR_n_* close to 1.0. Conversely, in low-confidence regions of SEQC2, a significant portion of reads are not haplotagged with either germline haplotype, leading to a notably lower *GHRR_n_* value. To ensure reliable variant calling, *V_t_*(*i*) is retained only if its *GHRR_n_* exceeds a predefined threshold 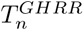

##### Germline Haplotype Read Count Filter

The analysis of high- and low-confidence variants in SEQC2 revealed that low-confidence variants predominantly exhibit low read counts (Supplementary Figure 16(A)). A naive approach is defining a minimum threshold of supporting reads for each variant 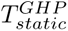, which cannot accommodate various degrees of tumor purity, heterogeneity, and aneuploidy. By investigating normal samples across multiple cell lines and tumor purity, we observed that high- and low-confidence variants often exhibit a bimodal distribution in their read count histograms (Supplementary Figure 16(B)), with two distinct peaks corresponding to high- and low-confidence variant clusters. We observed that high- and low-confidence variants form two distinct clusters (peaks) in read count histograms. The optimal threshold is the local minimum (valley) between these two peaks, which shifts based on the sample’s cell line and tumor purity. Our method automatically identifies this valley by smoothing the histogram, locating the two peaks, and finding the minimum between them (Supplementary Figure 17). The details of the three steps are described in the Supplementary Methods.

The LCVF filters reduce spurious variants and reshape the GHIR distribution, yielding a higher signal-to-noise ratio for subsequent purity estimation (Supplementary Figure 18).

#### 4.3.2 Purity estimation from the GHIR distribution

The median and IQR of the GHIR distribution are correlated with the tumor purity (Supplementary Figure 19). A second-order polynomial regression model for predicting tumor purity was trained using the median and IQR of the GHIR distributions using HCC1395_HKU, HCC1395_NYGC, COLO829_ONT, HCC1954_UCSC, HCC1937_UCSC, H2009_UCSC, and H1437_UCSC as training data and COLO829_NYGC as testing data. A range of tumor purity (0.2, 0.4, …, 1.0) was simulated by mixing the normal and tumor reads separately for each dataset. The model’s robustness was assessed using a leave-*k*-out cross-validation using the eight datasets.

### 4.4 Purity-aware somatic variant recalibration

We refine candidate somatic variants before final haplotagging using a multi-filter scheme guided by the estimated tumor purity (Supplementary Figure 20). First, a tumor-in-normal contamination (TINC) filter distinguish somatic variants contaminated in the matched normal from germline variants. Second, a read-consistency filter discards variants carrying reads with mixed haplotype alleles. Third, a haplottype-background filter removes variants with conflicting germline haplotypes. Finally, the variant-cluster filter removes clustered calls within dense intervals. For each filter, decision criteria are relaxed or tightened with respect to the estimated tumor purity. Candidates failing any filter are removed.

#### 4.4.1 TINC filter

The somatic variants can be present at low frequencies in the paired normal sample (Supplementary Figure 21(A)). Analysis of Variant Allele Frequencies (VAF) revealed that true somatic variants typically exhibit near-zero VAF in normal samples, while false variants demonstrate higher VAF and are likely to be putative germline variants (Supplementary Figure 21(B)). Because the frequency spectrum is correlated with the tumor purity, a filtering condition is applied to distinguish true TINC variants from germline variants. For each *V_t_*(*i*), if its VAF in the normal sample, *V AF_n_*(*i*), is less than a predefined low threshold 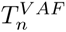, the variant is retained. Otherwise, it is discarded, where 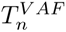 is dynamically determined based on the estimated tumor purity (see Section 4.4.5).

#### 4.4.2 Read-consistency filter

This filter examine the consistency of germline alleles of each read flanking a somatic variant. Each read suppointing a somatic variant is classified into one of four categories: read containing alleles consistent with only HP1 (*C_hp_*_1_(*i*)), read with alleles consistent with only HP2 (*C_hp_*_2_(*i*)), read containing alleles from both HP1 and HP2 (*C_mix_*(*i*)), and read without any germline alleles (*C_none_*(*i*)) (Supplementary Figure 22(A)). A Mixed Haplotype Read Ratio (*R_mix_*) for somatic variant at position *i* is defined as:

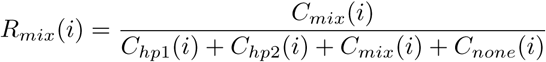

This ratio denotes the proportion of reads containing mixed alleles from both HP1 and HP2, relative to the total number of reads supporting *V_t_*(*i*). Analysis of the *R_mix_* distribution reveals that the majority of true somatic variants exhibit *R_mix_* values approaching zero (Supplementary Figure 22(B)), indicating these variants are predominantly supported by reads from a single germline haplotype. Conversely, the false variants often show *R_mix_* values near 1.0, suggesting they are supported by spurious reads containing alleles from both haplotypes. Based on this observation, a *V_t_*(*i*) is retained only if its *R_mix_*(*i*) is below a threshold *T_mix_* (Supplementary Figure 22(C)), which is dynamically adjusted based on the estimated tumor purity (see Section 4.4.5).

#### 4.4.3 Haplotagging-background filter

This filter examines the haplotype background of reads supporting each somatic variant. For each *V_t_*(*i*), the distribution of supporting reads across different haplotype backgrounds (HP1-1, HP2-1, and HP3, where HP3) is analyzed (Supplementary Figure 23(A)). Analysis of the haplotype backgrounds of supporting reads shows distinct distributions between true and false variants in three-dimensional scatter plots (Supplementary Figure 23(B)). In these plots, true somatic variants cluster along either axis, indicating predominant support from a single germline haplotype, while false variants appear scattered in the central region, suggesting support from multiple germline haplotypes. Let 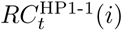 be reads derived from HP1 and 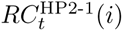 be reads derived from HP2 at *V_t_*(*i*) in tumor sample. A somatic variant *V_t_*(*i*) is discarded if both 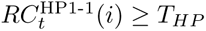 and 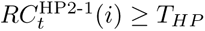 ; otherwise, *V_t_*(*i*) is retained, where *T_HP_* denotes the threshold for somatic read counts (Supplementary Figure 23 C) and is adaptively determined based on the estimated tumor purity (see Section 4.4.5).

#### 4.4.4 Variant cluster filter

We observed that some false somatic variants frequently co-occur on the same set of reads within a dense interval at low VAF (Supplementary Figure 24(A)). While true somatic variants may also arise within such regions, they typically appear on different sets of reads with distinct allele frequency patterns. A dense interval is defined as a genomic region in which any two variants separated by less than a predefined distance threshold (*T_dist_*, default 100 bp) (Supplementary Figure 24(B)). For each variant, *M_v_*(*i*) is defined as the average number of other nearby variants that appear on the same set of reads supporting the *i*-th variant (Supplementary Figure 24(C)). False variants generally exhibit higher *M_v_* values due to their co-occurrence on the same reads, while true somatic variants typically show lower *M_v_*values.

To distinguish true and false variants within these dense intervals, three criteria were required. First, since false variants typically exhibit lower VAF, we require both variant allele frequency (*V AF_t_*(*i*)) and alternative allele count (*ALT_t_*(*i*)) must be below thresholds *α* and *β* respectively. To ensure the interval contains a sufficient number of clustered variants, the total variant count (*SC_interval_*(*I*)) in the interval *I* must exceed a threshold *γ*. Finally, a Z-score (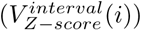) of *M_v_*(*i*) is calculated for each variant within each dense interval, with true variants typically appearing as false variants showing Z-scores near zero. A variant is discarded only when all these conditions are met: *V AF_t_*(*i*) *< α*, *ALT_t_*(*i*) *< β*, *SC_interval_*(*I*) *> γ*, and |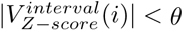, where the three thresholds are determined based on the estimated tumor purity (see Section 4.4.5).

#### 4.4.5 Purity-aware parameter optimization

The thresholds in the four filters (e.g., *V AF_t_*(*i*) *< α*, *ALT_t_*(*i*) *< β*) were adaptively optimized as a function of the estimated tumor purity. For each discrete purity level (e.g., 0.2, 0.4, …, 1.0), an independent parameter optimization was performed using simulation datasets derived from the six cell lines spanning a range of purity levels (Supplementary Figure 25). Optimization proceeded in two sequential phases. In the first phase, a global search was conducted by constrained random sampling across empirically defined parameter ranges to broadly explore the search space. In the second phase, a local refinement was performed using a grid search centered on the parameter regions yielding the highest performance during the global phase. The two-stage procedure produced a distinct set of optimal thresholds tailored to each tumor purity bin.

### 4.5 Integrated germline-somatic haplotype reconstruction

LongPhase-S consolidates outputs from the preceding modules to reconstruct germline and somatic haplotypes (Supplementary Figure 26). Similar to the preliminary haplotagging step, the germline and refined somatic variants are used to tag each read as germline (HP1 or HP2) or somatic haplotypes (HP1-1 or HP2-1), whereas reads without germline alleles are assigned to HP3 (e.g., a read too short to be phased with germline heterozygotes). To recover haplotype origin for HP3 reads, we implement a haplotype-inheritance procedure that leverages other reads carrying the same somatic allele yet with a consensus somatic haplotype background (Supplementary Figure 27). An HP3 read is assigned an inherited somatic haplotype only if all other reads are concordant for a single somatic background (i.e., either HP1-1 or HP2-1). Conversely, it is deemed non-inheritable if the supporting reads show somatic discordance or no reads are inheritable (e.g., all reads are HP3). Hence, LongPhase-S will tag each read as germline haplotypes (HP1, HP2) and somatic haplotypes (HP1-1, HP2-1, HP3).

### 4.6 Simulation of tumor-normal mixed purity

To systematically evaluate the performance of our method under varying tumor purity, we generated a series of *in silico* tumor-normal mixed samples. Starting with high-coverage pure tumor (100% purity) and matched normal BAM files, we created synthetic mixtures using the following three steps. First, the coverage of the input BAM files was determined using mosdepth (v0.3.3). Second, reads were randomly subsampled from the tumor and normal BAM files using “samtools view -s”. The sub-sampling fractions were chosen to achieve specific target purities while maintaining a fixed final tumor coverage of 50× and a normal coverage of 25×. Finally, the subsampled tumor and normal BAM files were merged using “samtools merge” to create the final mixed-purity BAM file. This procedure was repeated to generate datasets with tumor purities of 1.0, 0.8, 0.6, 0.4, and 0.2.

### 4.7 Data sources and read-level benchmarking

Eight publicly available tumor-normal paired ONT sequencing data were utilized for model training and evaluation, which include six cancer cell lines (HCC1395, COLO829, HCC1937, HCC1954, H1437, H2009) from multiple sequencing sources (Supplementary Table 1). For benchmarking somatic variant calling, the truth sets for HCC1395 were sourced from Sequencing Quality Control Phase 2 (SEQC2) [30], those for COLO829 were downloadedfrom New York Genome Center (NYGC) [31], and those for HCC1937, HCC1954, H1437, and H2009 were derived from Google DeepSomatic (orthogonal tools) [11]. The accuracy of somatic variant calling was computed using Illumina’s som.py (v0.3.12) [32],

We developed a somatic haplotagging benchmarking framework to quantify the accuracy of somatic haplotagging (Supplementary Figure 28). Becuase fully resolved somatic haplotypes are not available at this moment, we use the the well-validated somatic variants (SEQC2) of HCC1395 to produce a reference of somatic haplotagging reads. Specifically, each read carrying a somatic variant in the SEQC2 truth set was always assigned to one of the three somatic tags (HP1-1/HP2-1/HP3) based on the flanking germline variants. We then compared these SEQC2-tagged reads with those haplotagged by LongPhase-S to quantify somatic haplotagging accuracy at the read level. The haplotagged reads were classified as true positives if correctly assigned to their corresponding SEQC-2 tags, false positives if misassigned, and false negatives if unassigned. Performance was assessed using precision, recall, and F1-score. This framework thus provides an indirect, read-level evaluation of somatic haplotagging accuracy in the absence of fully resolved somatic haplotypes.

## 4.8 Code availability

The source code of LongPhase-S is freely available at https://github.com/CCU-Bioinformatics-Lab/longphase-s.

