## Supplementary Material for "LongPhase-S: purity estimation and variant recalibration with somatic haplotying for long-read sequencing"

#### Contents

**1 Supplementary Methods** **3**

**2 Supplementary Figures** **5**

**3 Supplementary Tables** **21**

### 1 Supplementary Methods

#### Bimodal Thresholding for Germline Haplotype Read Count Filter

We derive a filtering threshold from the inter-peak valley of a bimodal GHIR histogram. The procedure has three stages: (i) Gaussian smoothing of the raw histogram to reduce sampling noise while preserving modality; (ii) peak identification with adaptive height and distance de-duplication, followed by selection of representative (primary and secondary) peaks; (iii) inter-peak valley search with guardrails on cumulative mass and valley depth (Supplementary Figure 17). If the two peaks are not well-separated, a fixed minimum threshold is used.

##### Step 1: Gaussian Smoothing

Let $H[i]$ denote the raw GHIR histogram count at bin index $i \in \{0, \ldots, N_{\text{hist}} - 1\}$. We obtain a smoothed histogram $H_s[i]$ by convolving $H$ with a normalized, symmetric Gaussian kernel of width $\sigma$ (we explored $0.1 \le \sigma \le 2.0$). To keep the kernel symmetric, we use an odd kernel length

$$N_{\text{kernel}} = \begin{cases} \lfloor C_{\text{kernel}}\sigma + 1 \rfloor, & \text{if odd}, \\ \lfloor C_{\text{kernel}}\sigma + 1 \rfloor + 1, & \text{if even}, \end{cases}$$

where $C_{\text{kernel}}$ is a fixed multiplier (reported in code) and the center index is $k = \lfloor N_{\text{kernel}}/2 \rfloor$. The kernel entries are

$$K[j] = \frac{\exp\left(-\frac{1}{2}\left(\frac{j-k}{\sigma}\right)^2\right)}{\sum_{m=0}^{N_{\text{kernel}}-1} \exp\left(-\frac{1}{2}\left(\frac{m-k}{\sigma}\right)^2\right)}, \quad j = 0, \ldots, N_{\text{kernel}} - 1.$$

We use constant edge extension for indices outside $[0,\, N_{\text{hist}} - 1]$, i.e.,

$$\bar{H}(n) = \begin{cases} H[0], & n < 0, \\ H[n], & 0 \le n \le N_{\text{hist}} - 1, \\ H[N_{\text{hist}} - 1], & n \ge N_{\text{hist}}. \end{cases}$$

The smoothed histogram is then

$$H_s[i] = \sum_{j=0}^{N_{\text{kernel}}-1} K[j] \cdot \bar{H}(i + j - k).$$

##### Step 2: Peak Identification and Representative Peak Selection

Let $H_{\max} = \max_i H_s[i]$ be the maximum smoothed bin height. A bin $i$ is called a peak if it is a strict local maximum on $H_s$ (with boundary handling) and exceeds an adaptive threshold

$$T_{\text{peak}} = \max\left(r_{\min} \cdot H_{\max},\ t_{\min}\right),$$

where the minimum relative peak ratio $r_{\min} \in [0.02, 0.10]$ and minimum absolute count $t_{\min} \in [5, 20]$ (cohort-dependent) were explored. To avoid duplicates, peaks closer than a minimum separation $d_{\min} \in [2, 5]$ bins are merged by retaining the higher one.

We index detected peaks in ascending bin order as $\text{peak}_0, \text{peak}_1, \ldots$. For two consecutive peaks $\text{peak}_i$ and $\text{peak}_{i+1}$, define

$$\text{trend}(i \to i+1) = \begin{cases} \text{UP}, & H_s[\text{peak}_i] < H_s[\text{peak}_{i+1}], \\ \text{DOWN}, & H_s[\text{peak}_i] > H_s[\text{peak}_{i+1}], \\ \text{FLAT}, & H_s[\text{peak}_i] = H_s[\text{peak}_{i+1}]. \end{cases}$$

We label as *main peaks* those at up–down transitions; for edges, we retain the first peak if its rightward trend is DOWN and the last peak if its leftward trend is UP. Let $\text{peak}_{h1}$ and $\text{peak}_{h2}$ be the tallest and second-tallest peaks by height $H_s[\cdot]$. If there is only one main peak, that peak is the *primary* peak $\text{peak}_{\text{prim}}$. Otherwise, set $\text{peak}_{\text{prim}}$ to the rightmost of $\{\text{peak}_{h1}, \text{peak}_{h2}\}$ (tie-break by index). To choose the *secondary* peak $\text{peak}_{\text{sec}}$, scan leftward from $\text{peak}_{\text{prim}}$. If $\text{peak}_{\text{prim}}$ is the second peak, take its immediate left neighbor. Otherwise, select the nearest peak that sits at a DOWN–UP transition; if none exists, fall back to the leftmost peak.

##### Step 3: Inter-Peak Valley Search and Threshold

Let $i_{\text{sec}}$ be the index of $\text{peak}_{\text{sec}}$, and let $i_{\text{prev}}$ and $i_{\text{next}}$ be the indices of the nearest *main* peaks to its left and right, respectively. A candidate valley is any strict local minimum $i$ satisfying

$$H_s[i] < H_s[i-1] \quad \text{and} \quad H_s[i] < H_s[i+1].$$

Define the right and left candidate intervals

$$\mathcal{L}_{\text{next}} = \{\, i \mid i_{\text{sec}} < i < i_{\text{next}} \,\}, \qquad \mathcal{L}_{\text{prev}} = \{\, i \mid i_{\text{prev}} < i < i_{\text{sec}} \,\}.$$

For any index $i$, define the cumulative mass $cd[i] = \sum_{n=0}^{i} H_s[n]$ and total mass $cd_{\text{tot}} = \sum_{n=0}^{N_{\text{hist}}-1} H_s[n]$, with proportion $p_{\text{valley}} = cd[i]/cd_{\text{tot}}$. We first search $\mathcal{L}_{\text{next}}$ and accept a candidate there only if $p_{\text{valley}} < p_{\text{limit}}$ with $p_{\text{limit}} \in [0.6, 0.8]$; otherwise we fall back to $\mathcal{L}_{\text{prev}}$. In either interval, choose the deepest valley (minimal $H_s[i]$); denote its index by $i_{\text{valley}}$ and depth by $h_{\text{valley}} = H_s[i_{\text{valley}}]$. With $h_{\text{limit}} \in [0.4, 0.6]$ the maximum allowable relative valley height, the bimodal valleythreshold is

$$T_{\text{Bimodal}} = \begin{cases} i_{\text{valley}}, & \text{if } h_{\text{valley}} \le H_{\max} \cdot h_{\text{limit}}, \\ 0, & \text{otherwise}, \end{cases}$$

where $H_{\max} = \max_i H_s[i]$.

### 2 Supplementary Figures

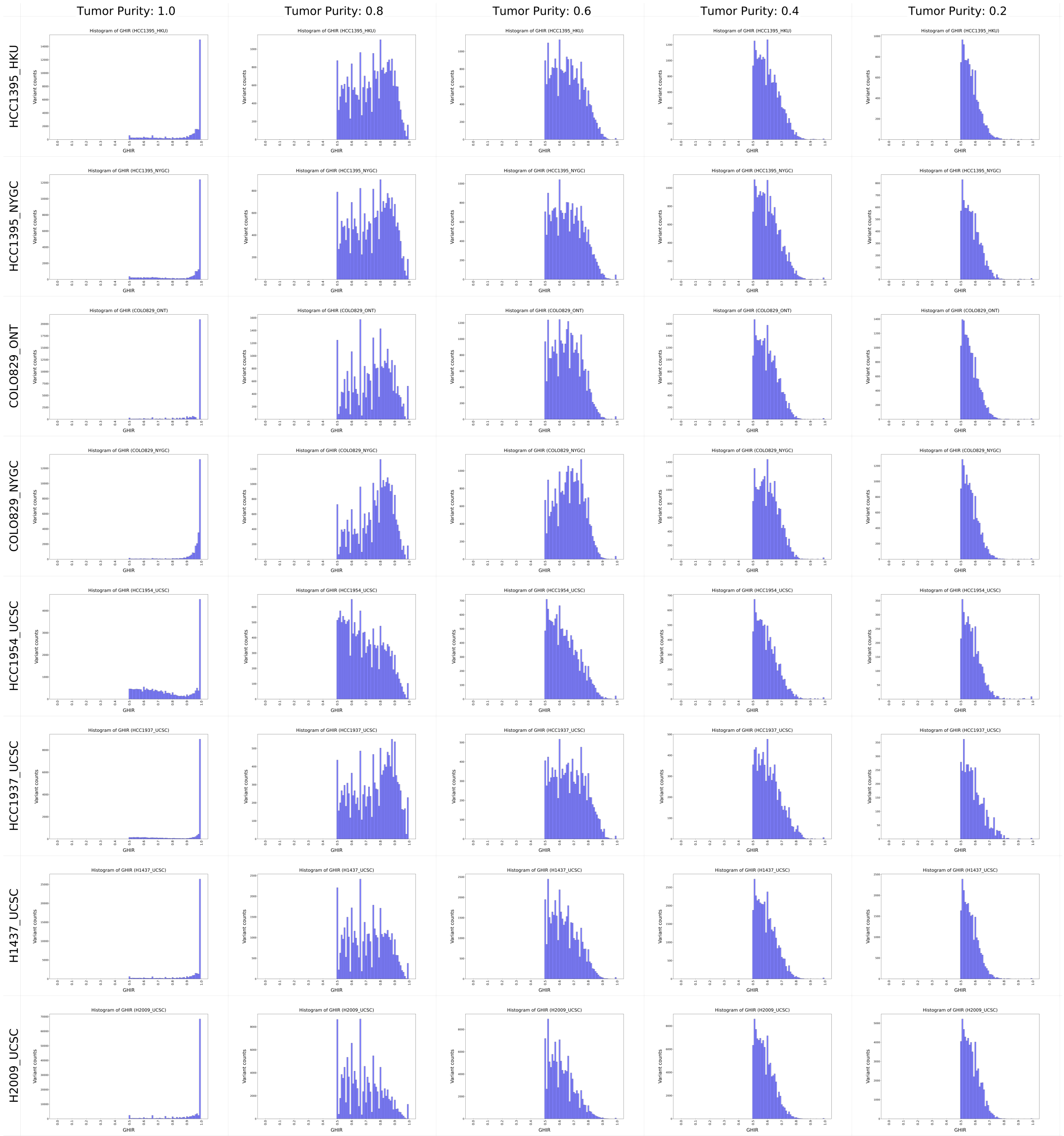

**Supplementary Figure 1**: The GHIR distributions of eight cancer cell line datasets at varying purity levels (1.0 to 0.2) after LCVF. The systematic changes in distribution patterns correlating with tumor purity levels suggest that GHIR median and dispersion are relevant to purity estimation

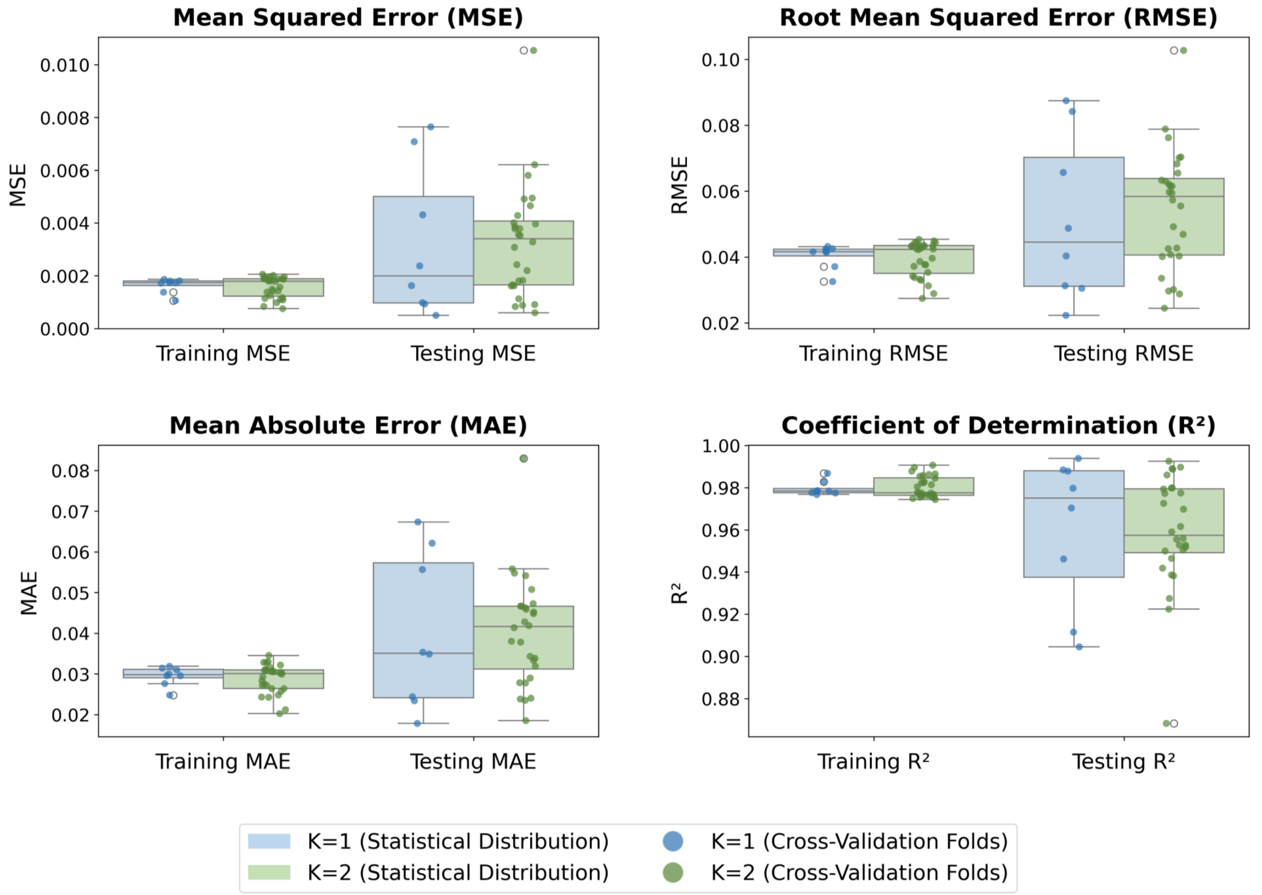

**Supplementary Figure 2**: Distribution of performance metrics from Leave-k-Out cross-validation. The box plots show the distribution of MSE, RMSE, MAE, and $R^2$ for both training and testing sets under $k = 1$ and $k = 2$, with similar distributions observed across all metrics.

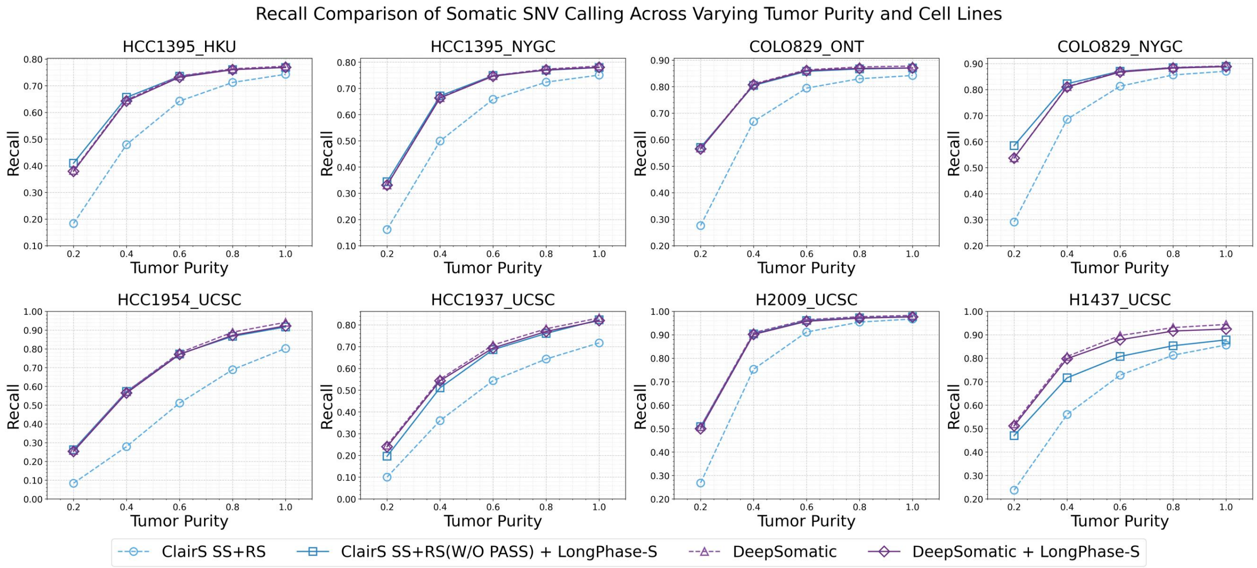

**Supplementary Figure 3**: Recall of four somatic SNV calling pipelines across varying tumor purity levels in synthetic tumor datasets.

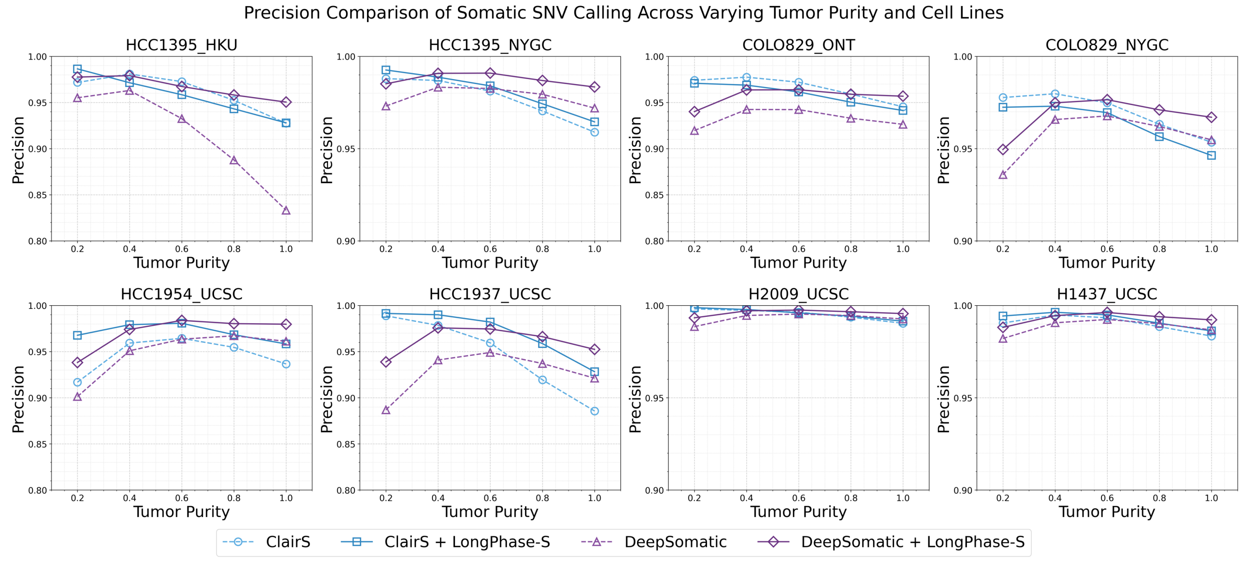

**Supplementary Figure 4**: Precision of four somatic SNV calling pipelines across varying tumor purity levels in synthetic tumor datasets.

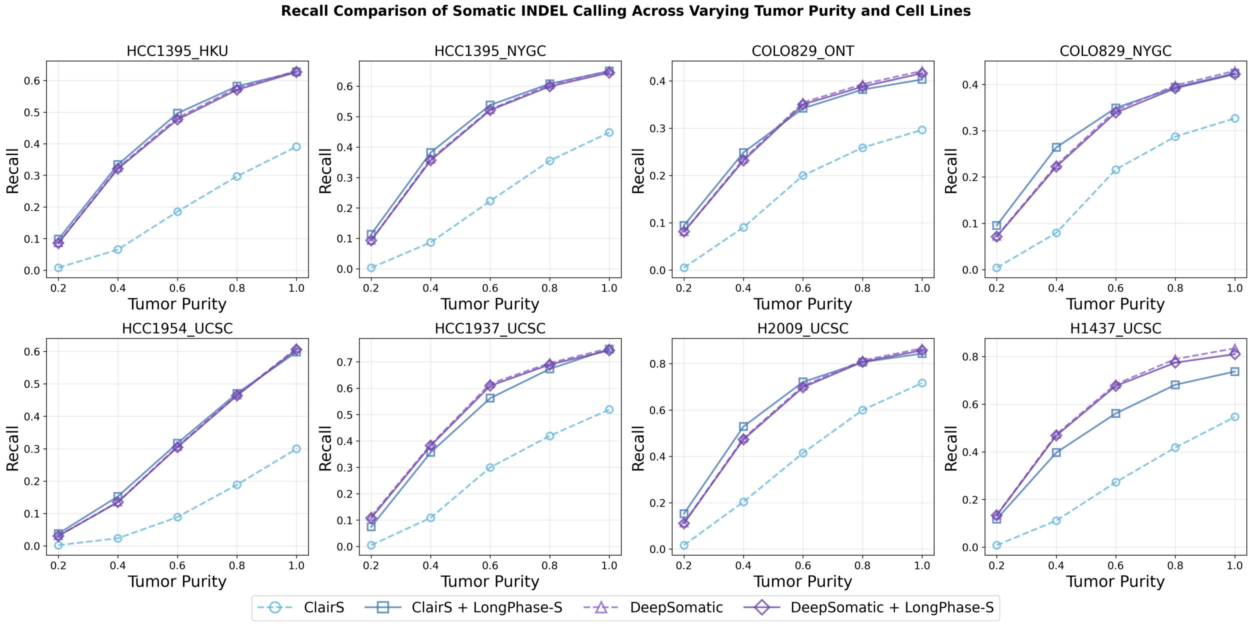

**Supplementary Figure 5**: Recall of four somatic indel calling pipelines across varying tumor purity levels in synthetic tumor datasets.

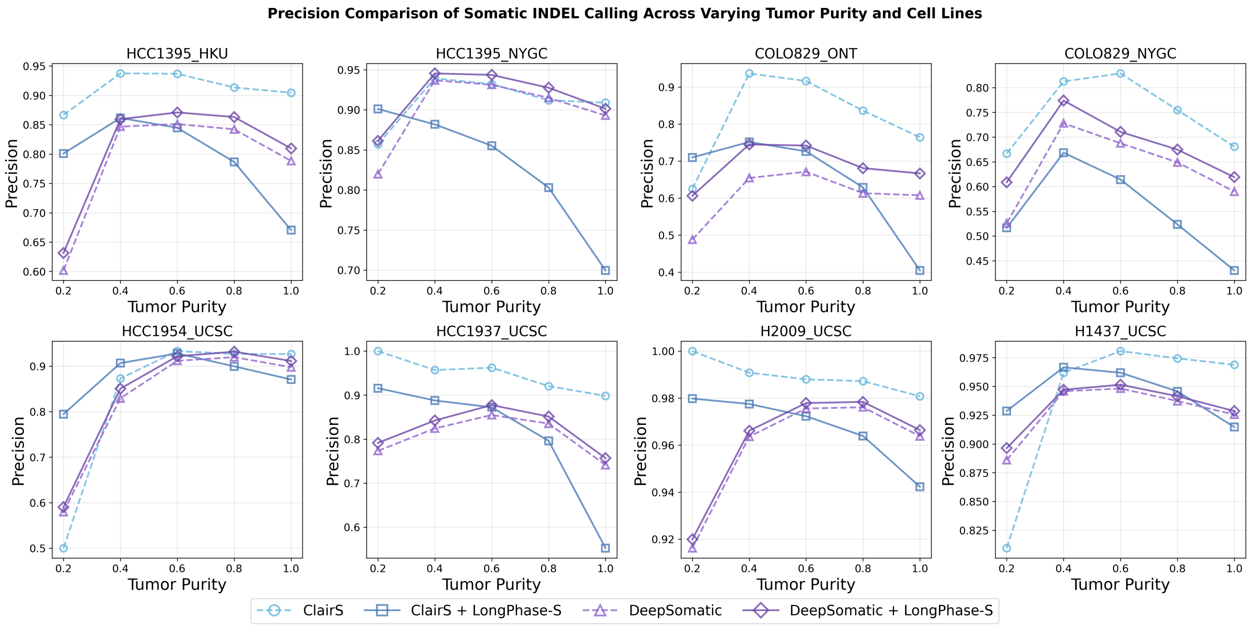

**Supplementary Figure 6**: Precision of four somatic indel calling pipelines across varying tumor purity levels in synthetic tumor datasets.

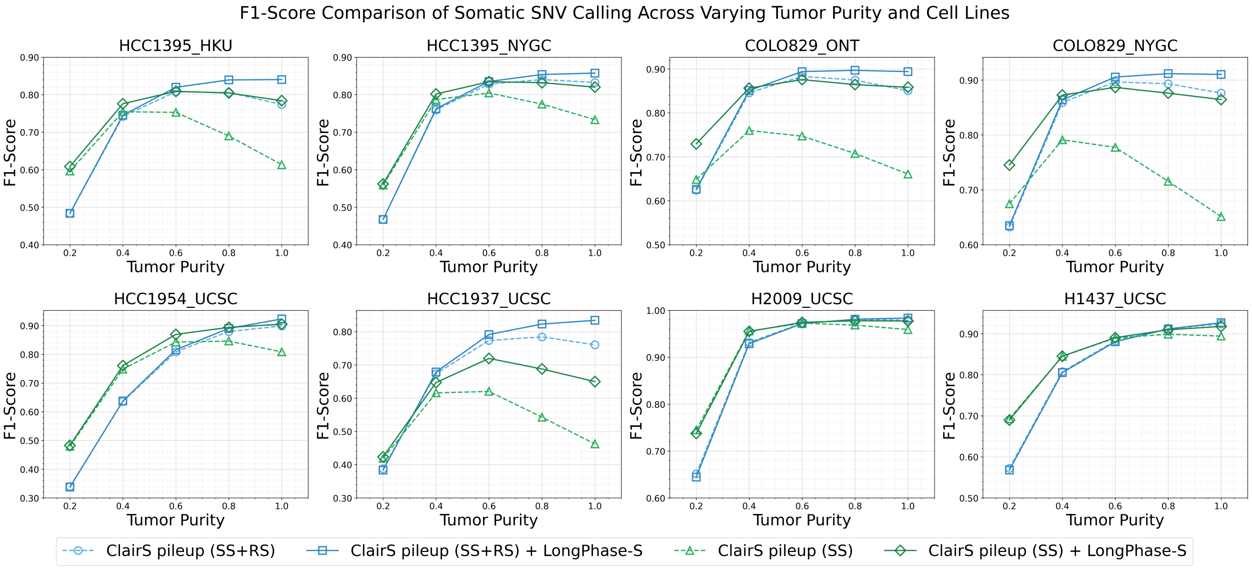

**Supplementary Figure 7**: F1-Score of the ClairS pileup model with and without LongPhase-S across synthetic tumor purities.

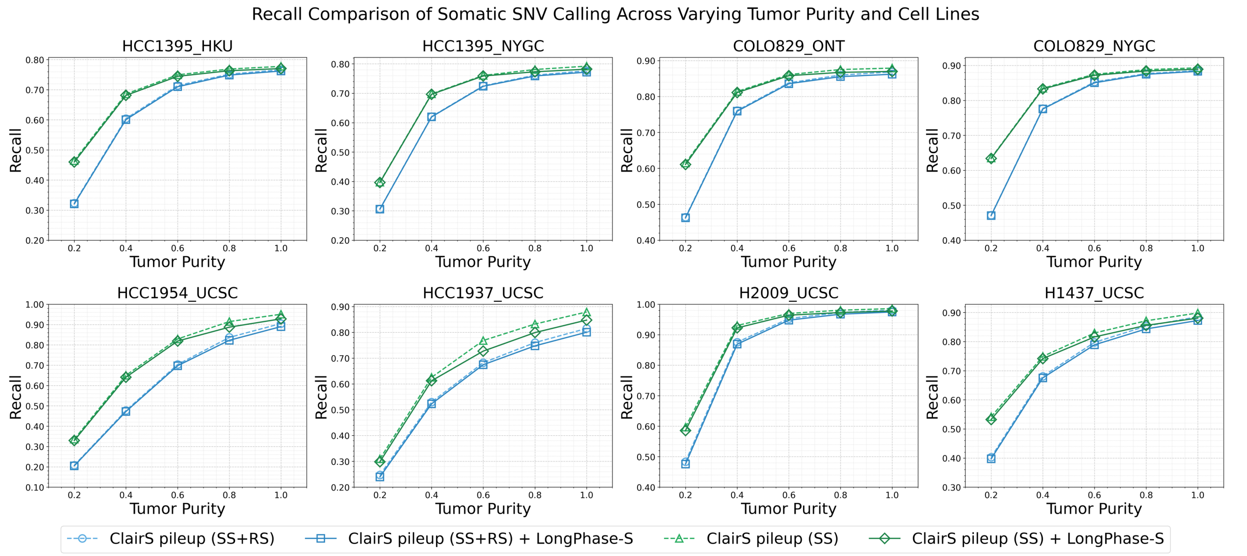

**Supplementary Figure 8**: Recall of the ClairS pileup model with and without LongPhase-S across synthetic tumor purities.

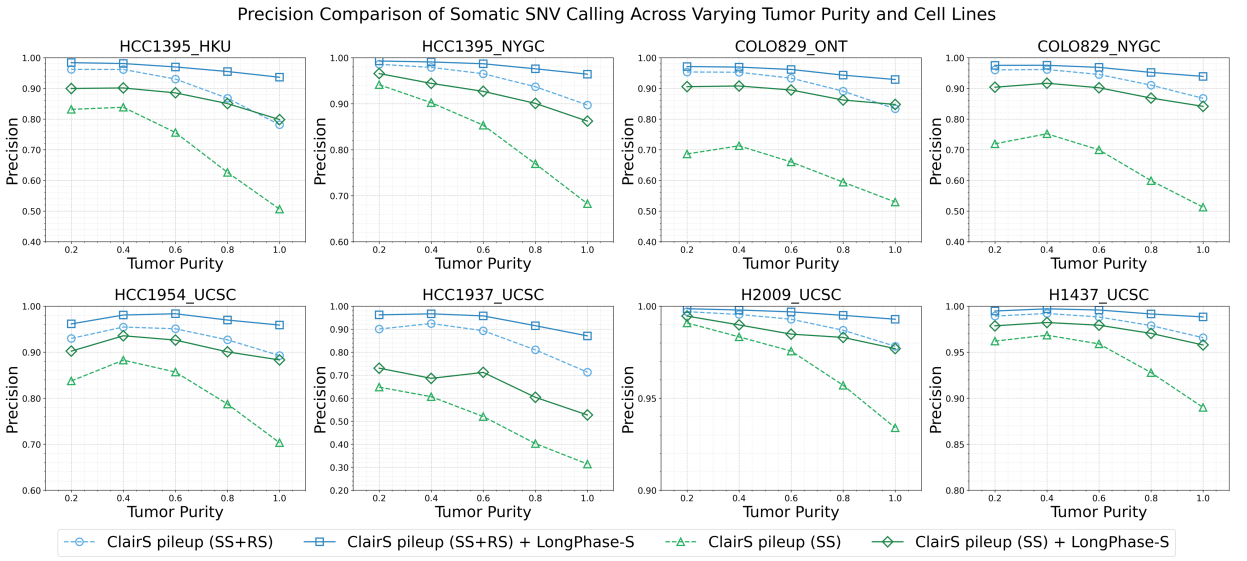

**Supplementary Figure 9**: Precision of the ClairS pileup model with and without LongPhase-S across synthetic tumor purities.

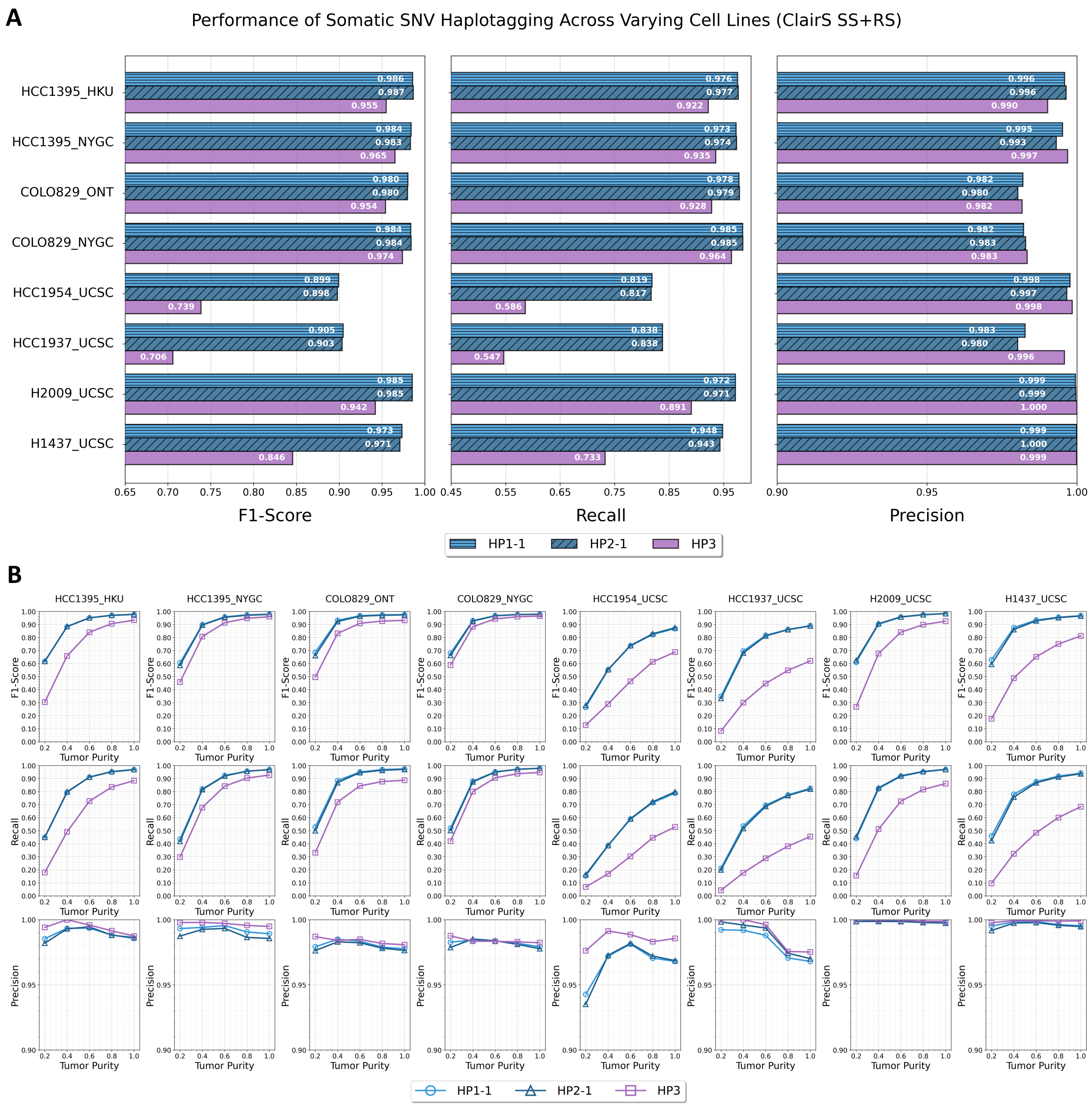

**Supplementary Figure 10**: Read-level accuracy of somatic haplotagging (HP1-1, HP2-1, and HP3) by ClairS (SS+RS model) across (a) eight cancer cell line datasets and (B) varying tumor purities.

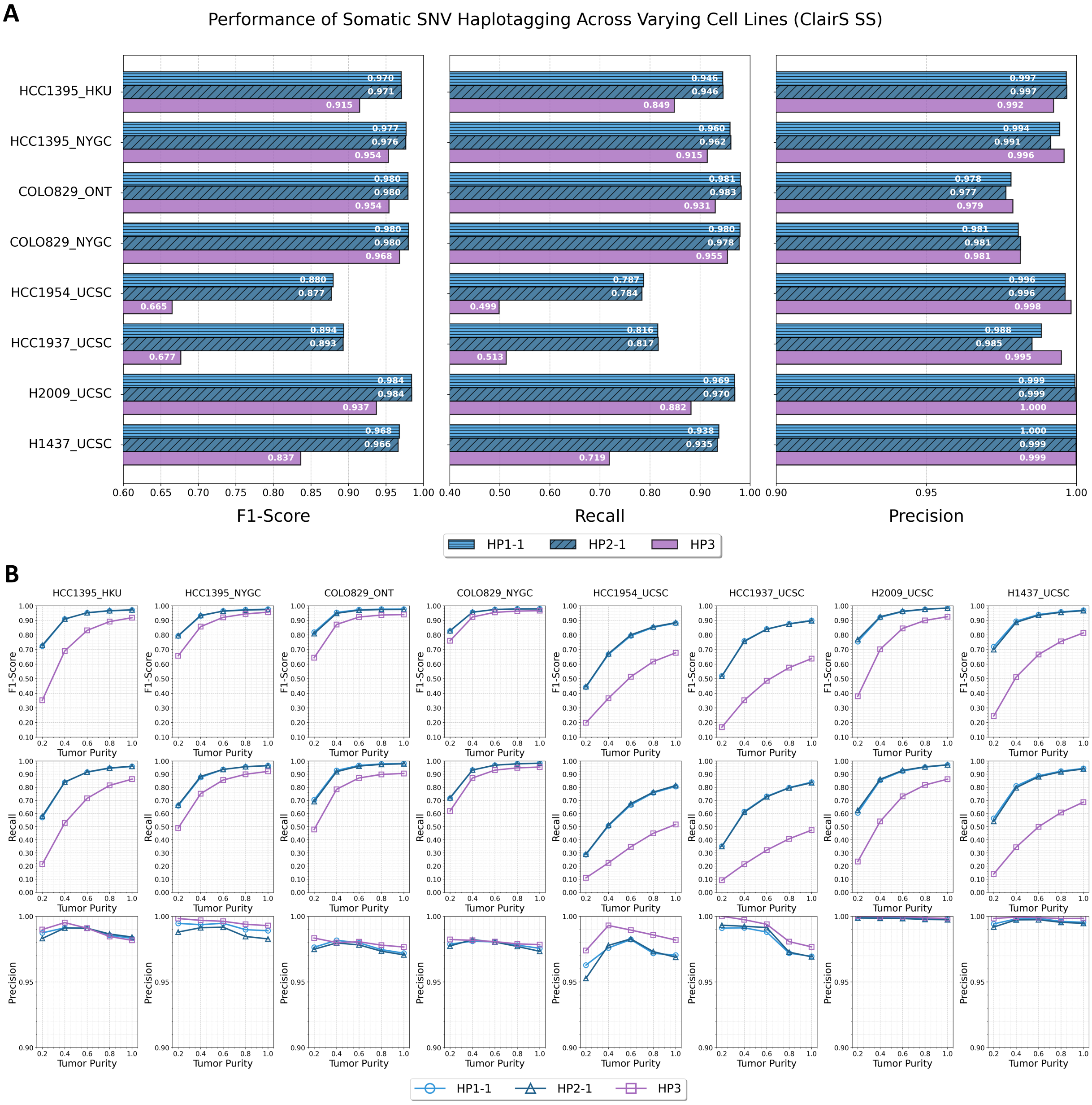

**Supplementary Figure 11**: Read-level accuracy of somatic haplotagging (HP1-1, HP2-1, and HP3) by ClairS (SS model) across (a) eight cancer cell line datasets and (B) varying tumor purities.

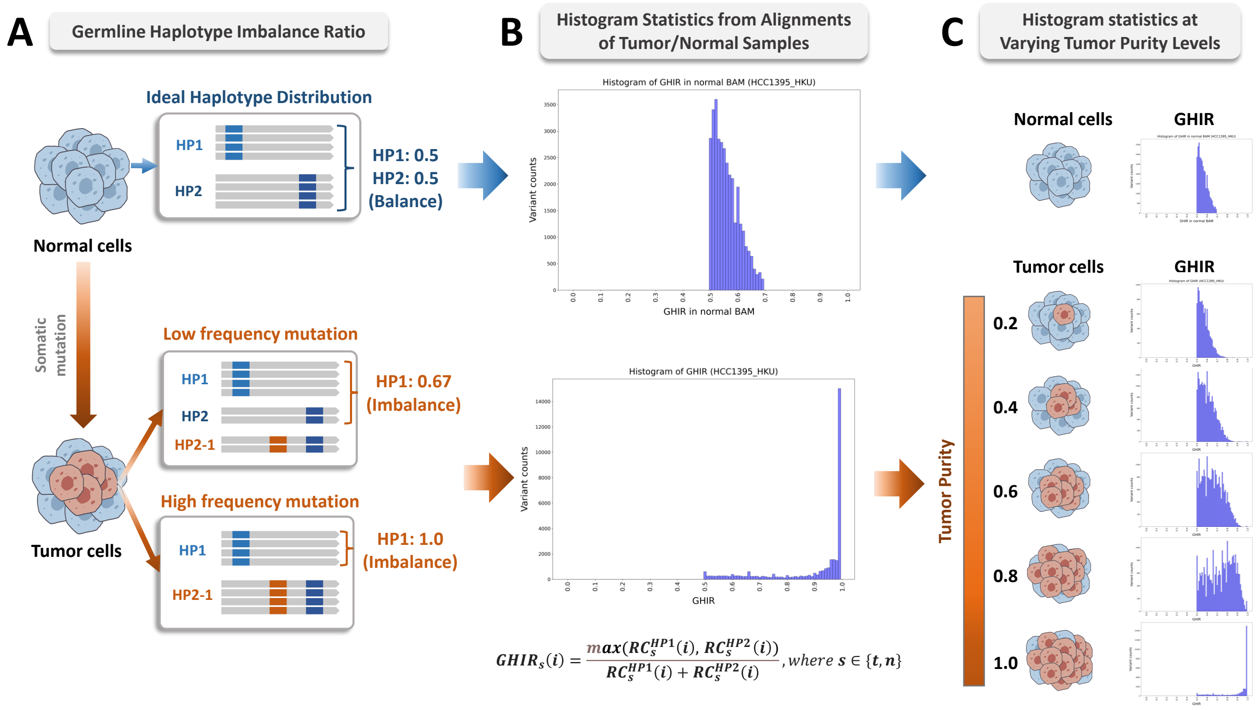

**Supplementary Figure 12**: Illustration of haplotype imbalance induced during tumorigenesis using GHIR. (A) Conceptual diagram depicting how somatic mutations affect germline haplotype balance. (B) GHIR distribution in alignments of normal and tumor samples from the HCC1395_HKU cell line. (C) GHIR distributions across different tumor purity levels in synthetic tumor samples.

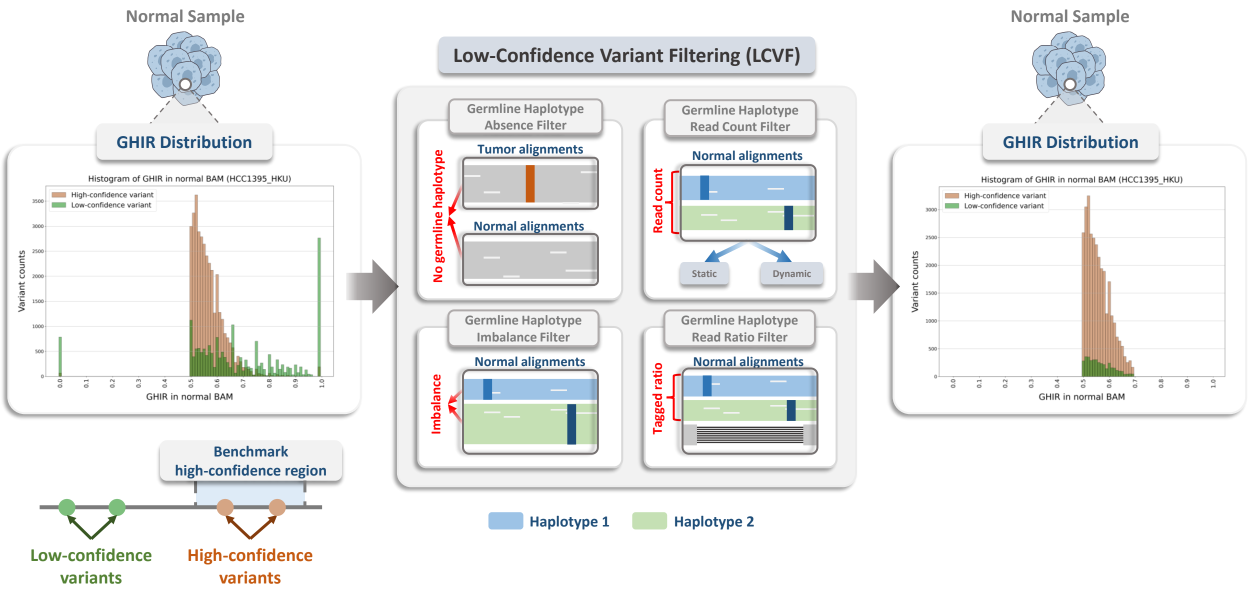

**Supplementary Figure 13**: Workflow of the Low-Confidence Variant Filtering.

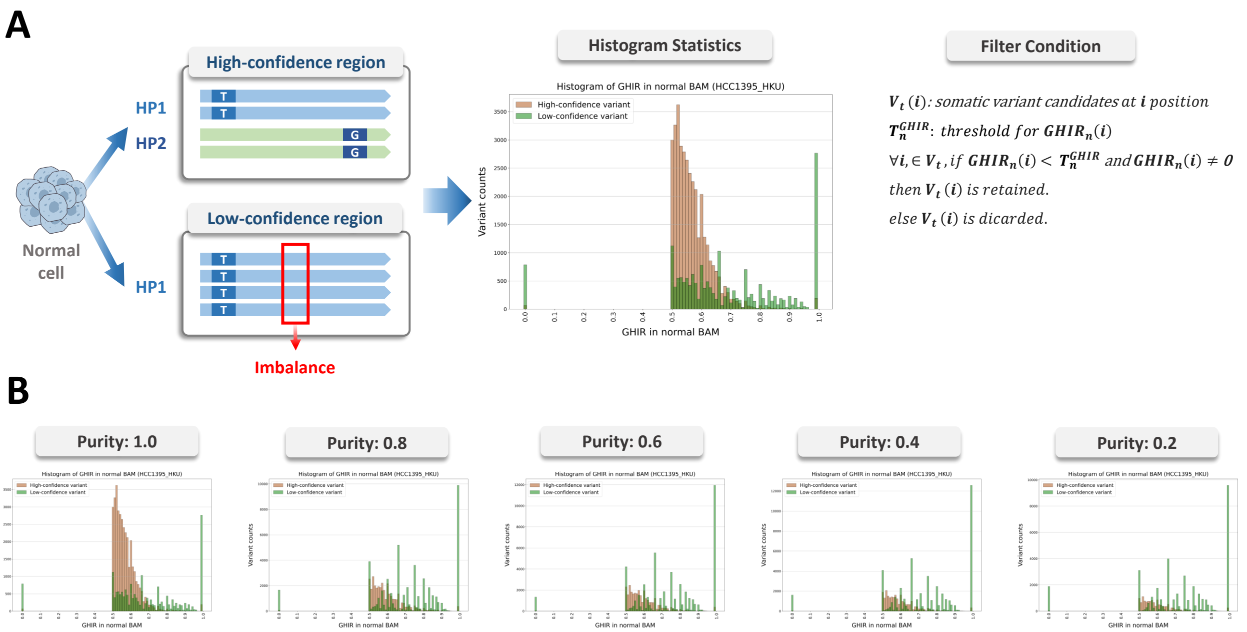

**Supplementary Figure 14**: Schematic of the Germline Haplotype Imbalance Filter. (A) Conceptual diagram showing how GHIR is used to identify high-confidence and low-confidence regions in normal sample alignments. (B) GHIR distribution in normal sample of HCC1395_HKU cell line showing the separation of high-confidence variants (brown) and low-confidence variants (green), where high-confidence variants cluster around GHIR = 0.5.

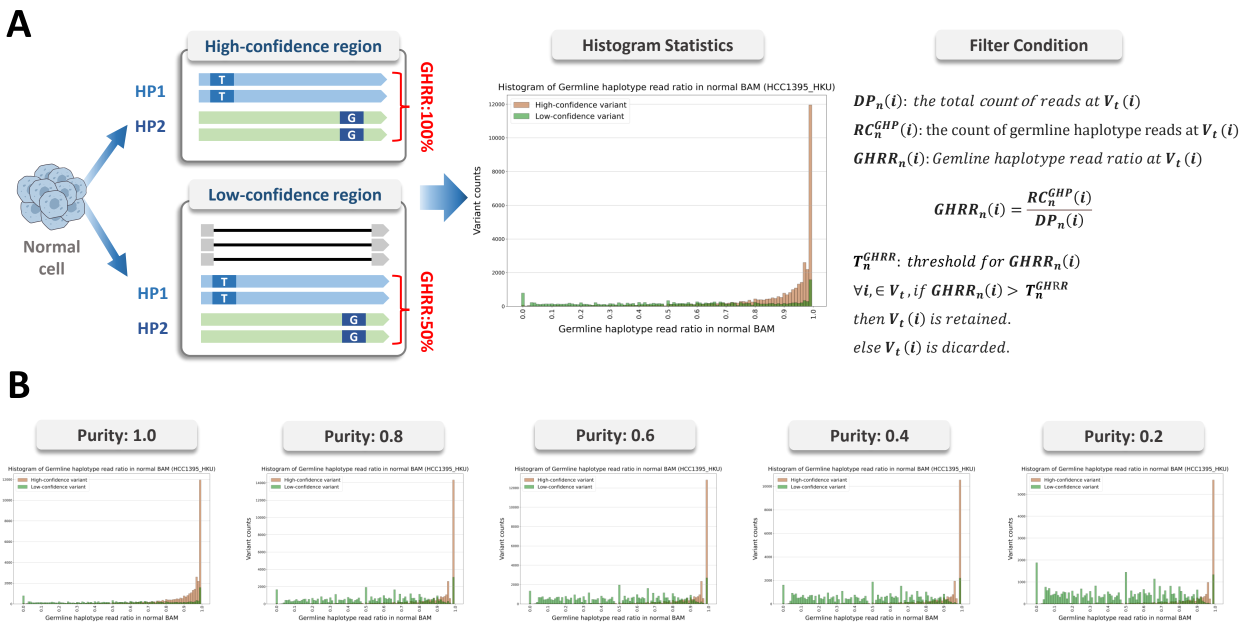

**Supplementary Figure 15**: Schematic of the Germline Haplotype Read Ratio Filter. (A) Conceptual diagram showing how GHRR distinguishes between high-confidence and low-confidence regions based on read support for germline haplotypes. (B) GHRR distribution in normal sample of HCC1395_HKU cell line, where high-confidence variants (brown) show GHRR values close to 1.0, indicating strong support for germline haplotypes.

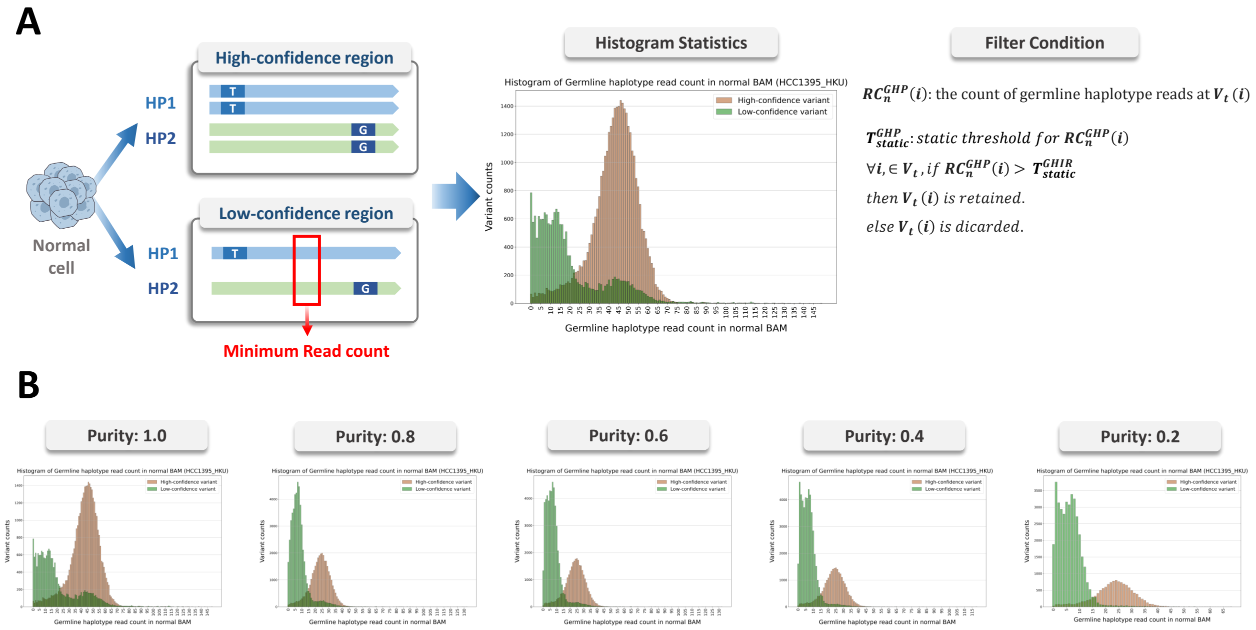

**Supplementary Figure 16**: Schematic of the Germline Haplotype Read Count Filter (static approach). (A) Conceptual diagram showing how read count thresholds distinguish between high-confidence and low-confidence regions in normal sample alignments. (B) Distribution of germline haplotype read counts in HCC1395_HKU cell line at varying tumor purity levels.

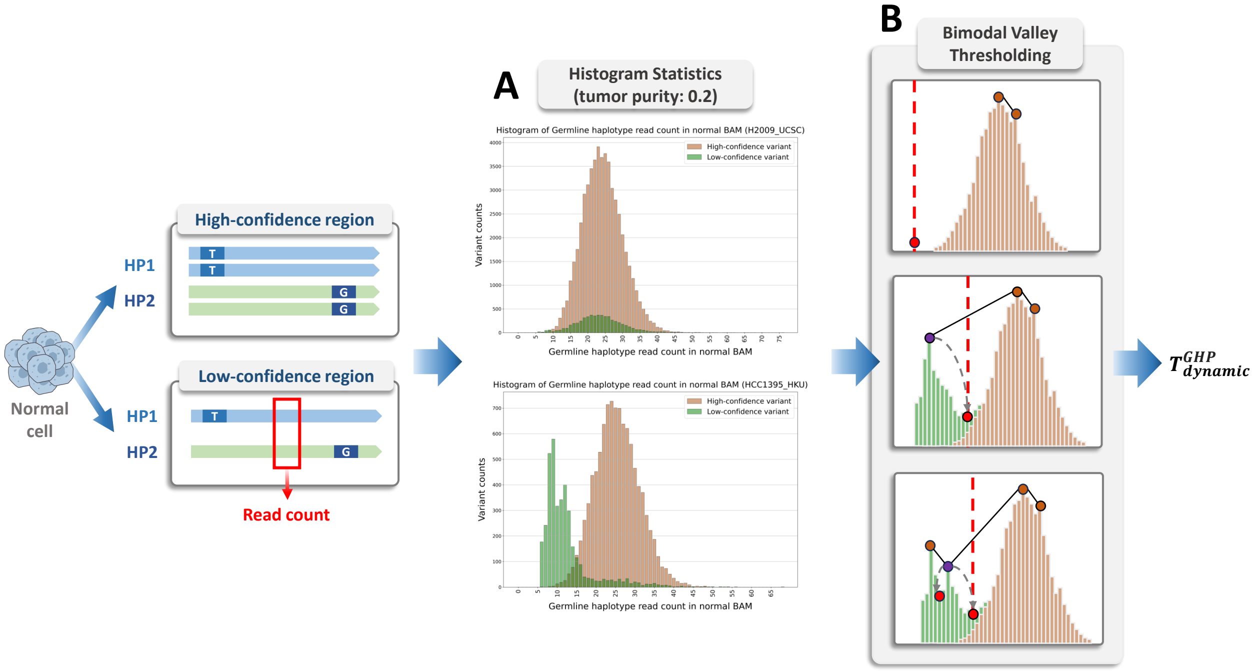

**Supplementary Figure 17**: Schematic of the Bimodal Valley Thresholding. (A) Histogram statistics of germline haplotype read counts in normal samples from two different cell lines (H2009_UCSC and HCC1395_HKU), showing the distribution of high- and low-confidence variants. (B) The process of Bimodal Valley Thresholding, demonstrating the identification of valley points between high-confidence and low-confidence regions to determine the dynamic threshold $T_{dynamic}^{GHP}$.

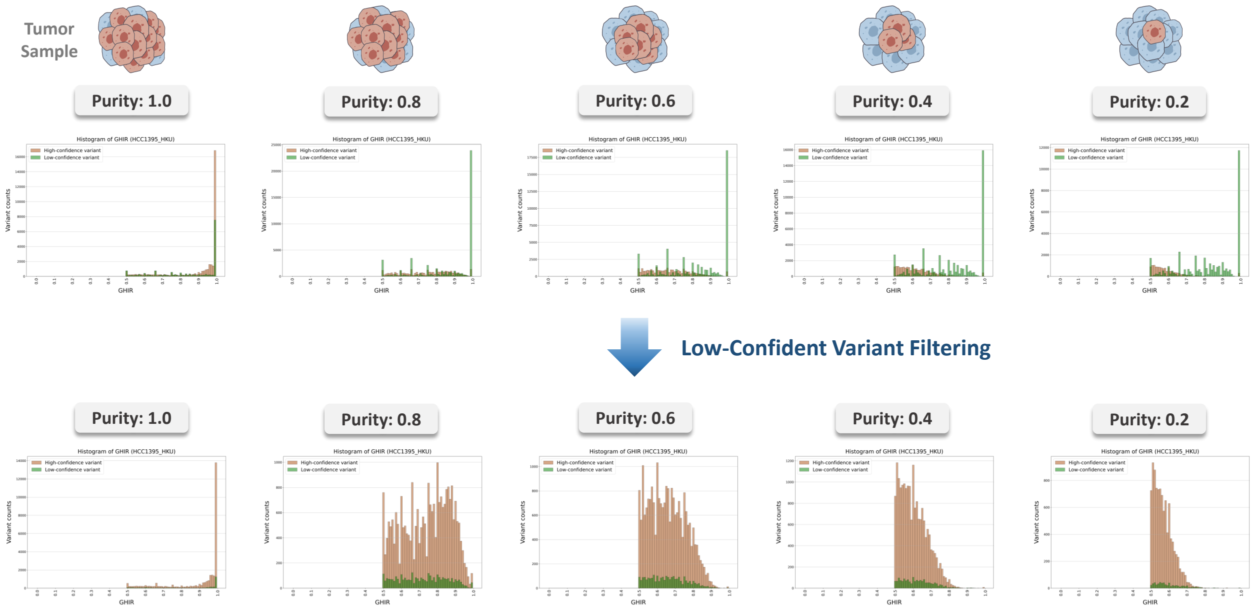

**Supplementary Figure 18**: A comparative visualization showing histograms of the GHIR in HCC1395_HKU synthetic tumor sample before and after the application of LCVF, demonstrating that the filtering process effectively removes noise and clarifies the signal related to tumor purity.

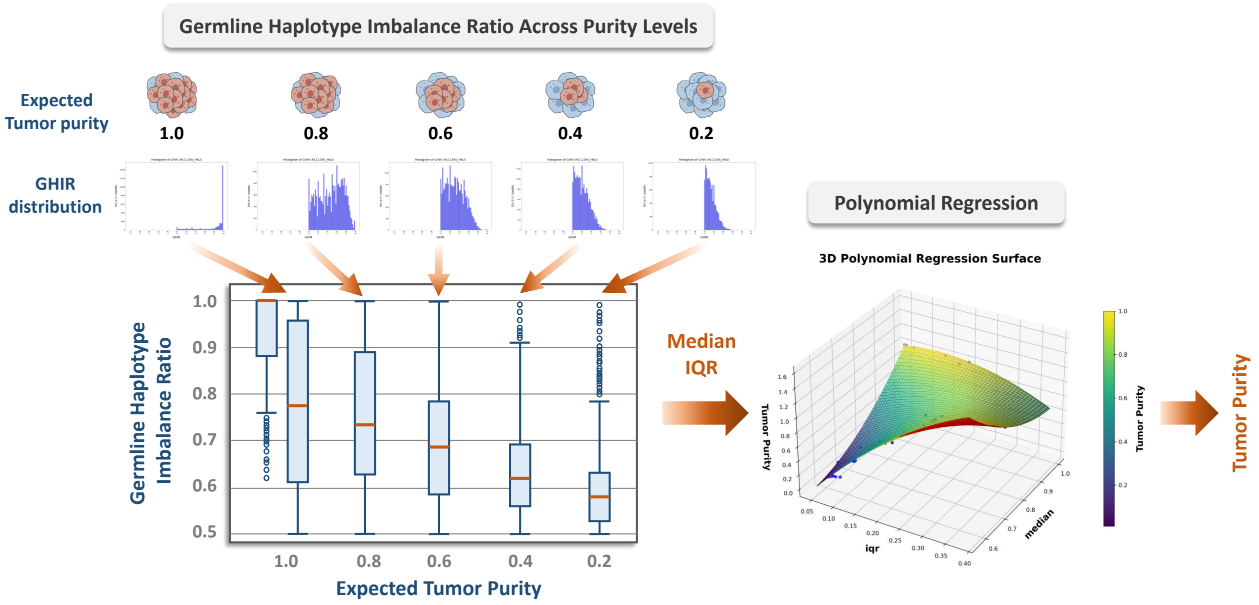

**Supplementary Figure 19**: Statistical features (median and IQR) extraction from GHIR distributions and their use in training a 2-degree polynomial regression model for tumor purity estimation.

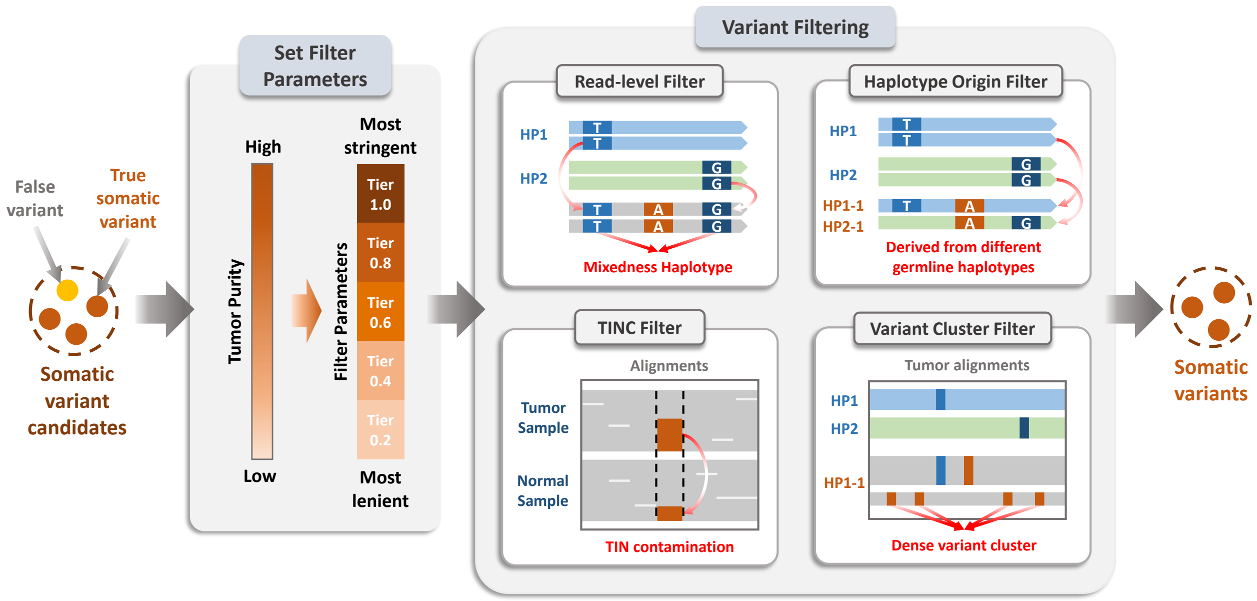

**Supplementary Figure 20**: Workflow of the tumor purity-aware somatic variant calling, which uses an estimated tumor purity value to dynamically set parameters for four parallel filters designed to remove different classes of false-positive variants.

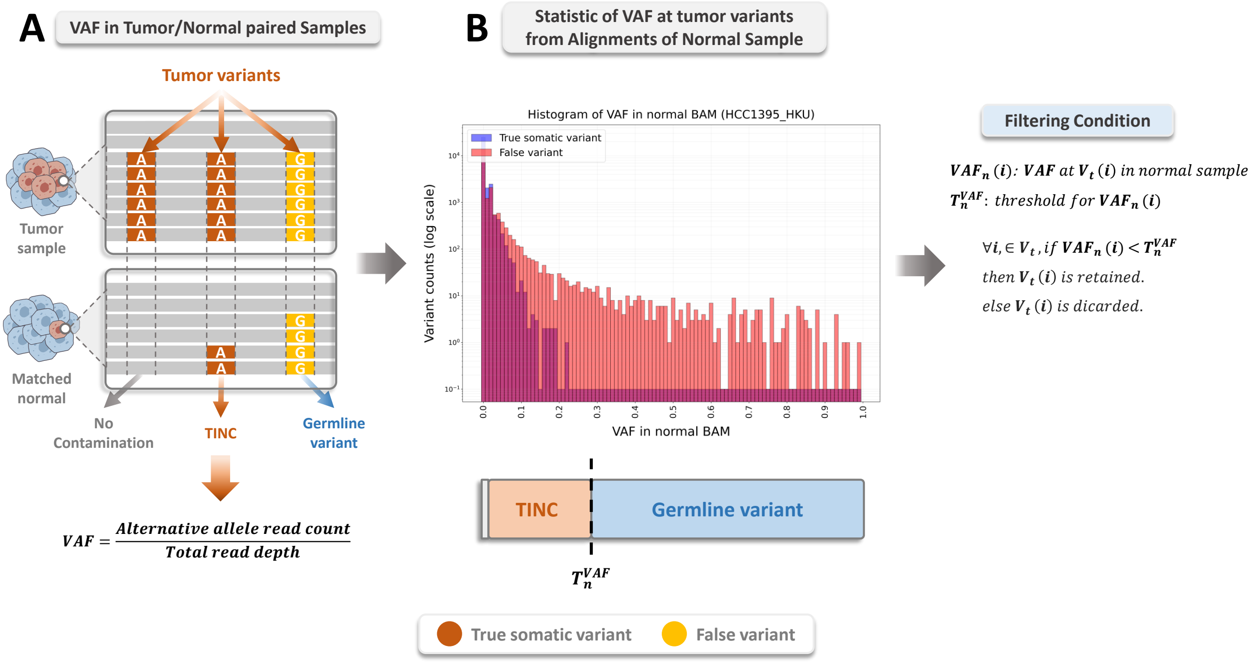

**Supplementary Figure 21**: Schematic of the TINC filter. (A) Conceptual diagram comparing tumor variants with their corresponding signals in the matched normal sample. (B) Histogram of VAF in the normal sample of the HCC1395_HKU cell line. True somatic variants tend to have a VAF near zero, while false variants exhibit a range of non-zero VAFs. This difference in VAF distribution is used to filter candidate somatic variants by applying a low VAF threshold.

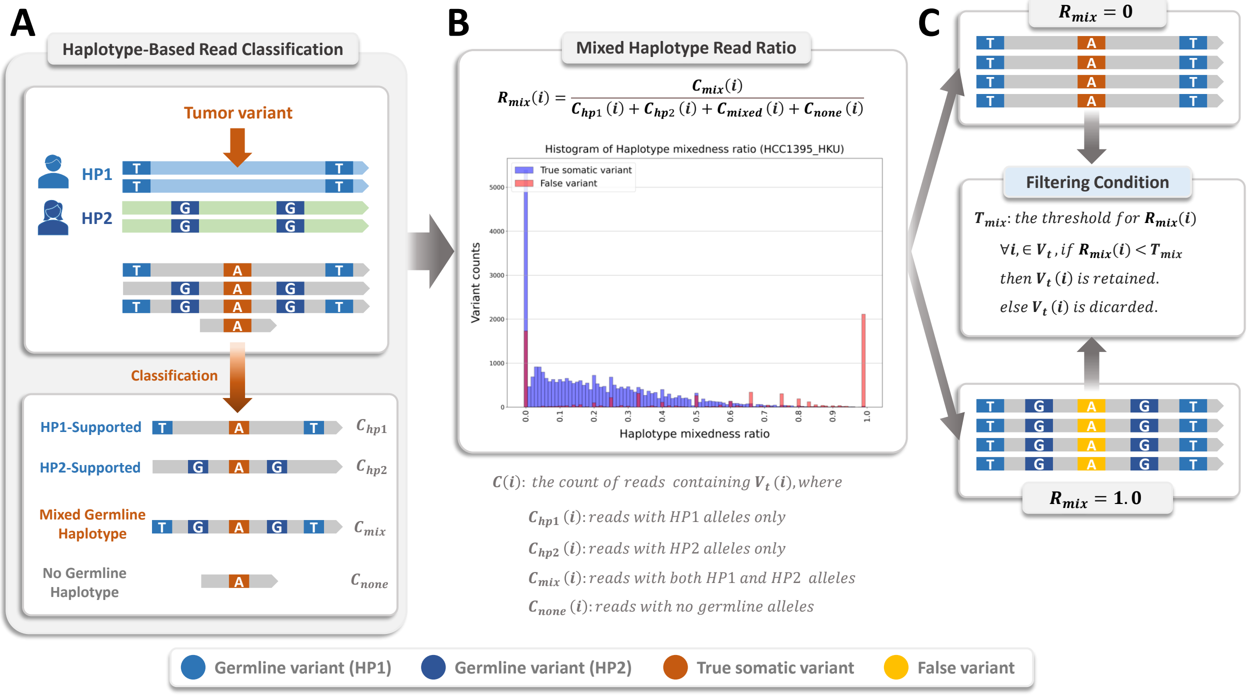

**Supplementary Figure 22**: Schematic of the read-level filter. (A) Classification of tumor reads according to germline haplotype context (HP1-supported, HP2-supported, mixed, or none). (B) Distribution of $R_{mix}$ in the HCC1395_HKU cell line, showing true somatic variants (blue) clustering near 0 and a subset of false variants (red) enriched near 1.0. (C) Filtering condition based on $R_{mix}$, where candidate variants with $R_{mix}(i)$ values exceeding the threshold are discarded.

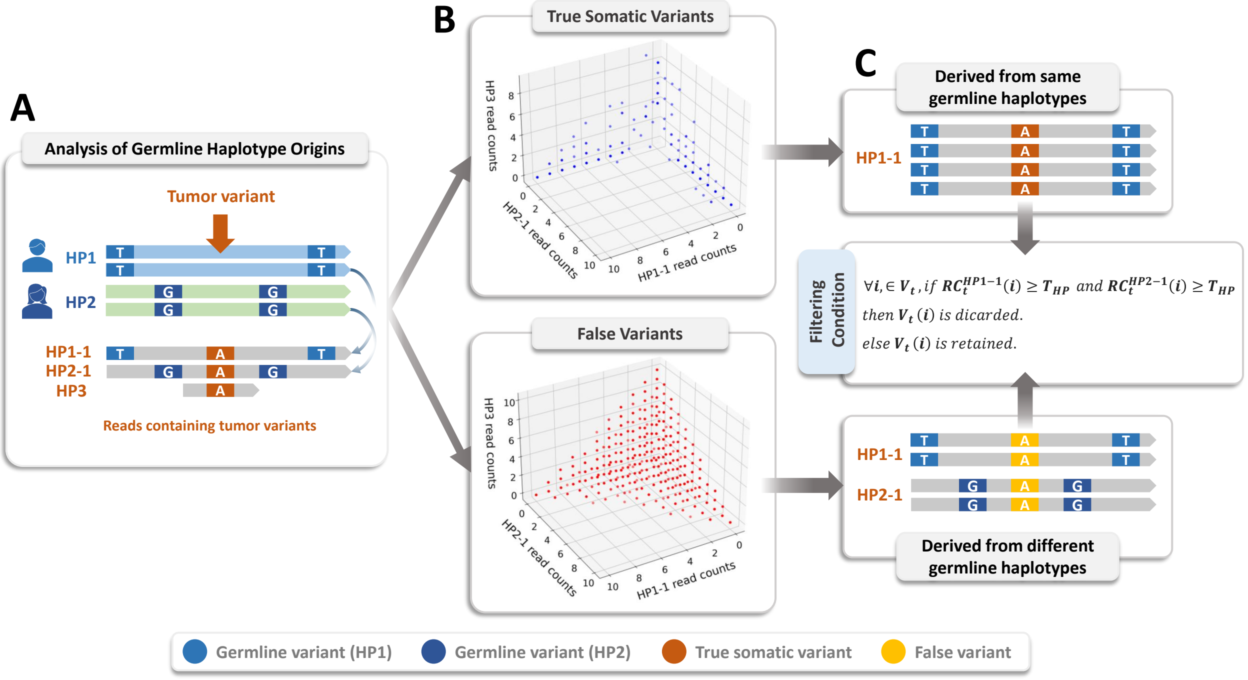

**Supplementary Figure 23**: Schematic of haplotype origin filter. (A) Analysis of germline haplotype origins, where tumor variant-containing reads are assigned to somatic haplotypes. (B) Three-dimensional scatter plots of low-VAF variants in the HCC1395_HKU cell line show true somatic variants (blue) clustering along the axes, supported by a single haplotype, whereas false variants (red) scatter centrally, reflecting mixed haplotype contributions and artifacts. (C) Filtering condition: a $V_t(i)$ is discarded if reads supporting it are observed above the threshold in both HP1-1 and HP2-1; otherwise, the variant is retained.

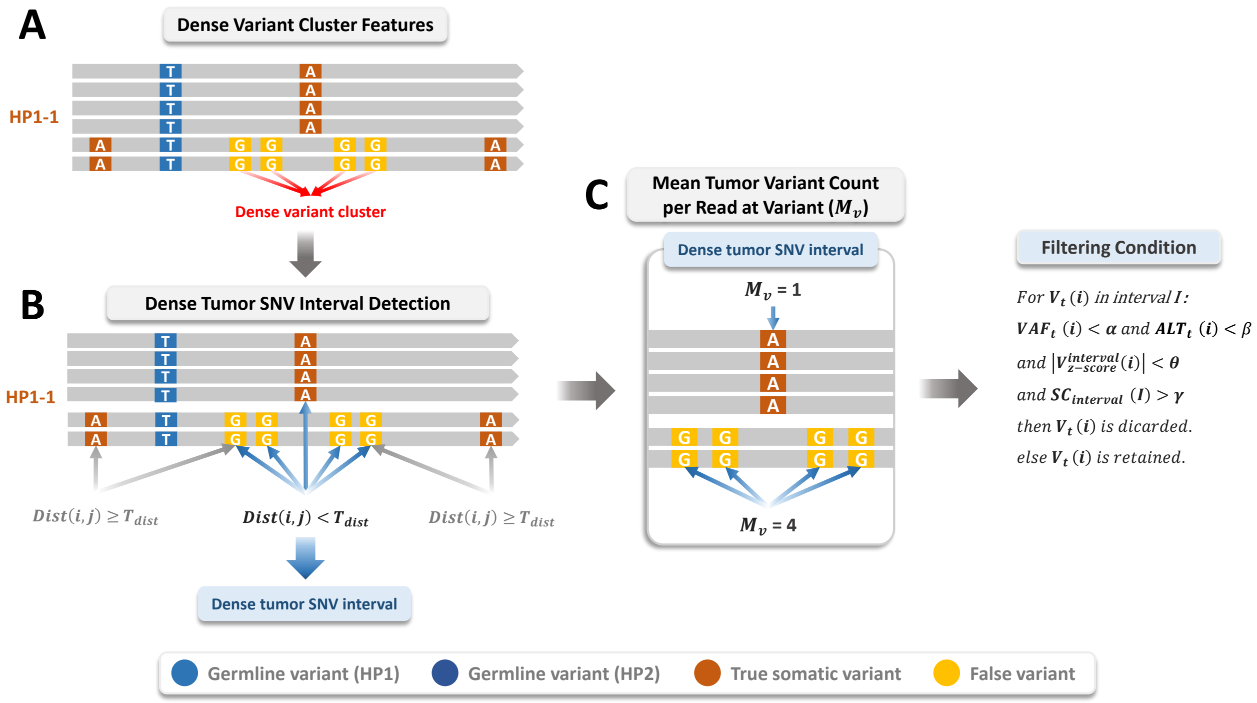

**Supplementary Figure 24**: Schematic of the variant cluster filter. (A) A simple example of dense variant cluster features showing true somatic variants and clustered false variants on reads from HP1-1. (B) Dense tumor SNV interval detection based on genomic distance ($T_{dist}$) between variants. (C) Mean variant count per read ($M_v$) analysis within dense intervals, demonstrating the difference between true variants ($M_v = 1$) and false variants ($M_v = 4$) in this example.

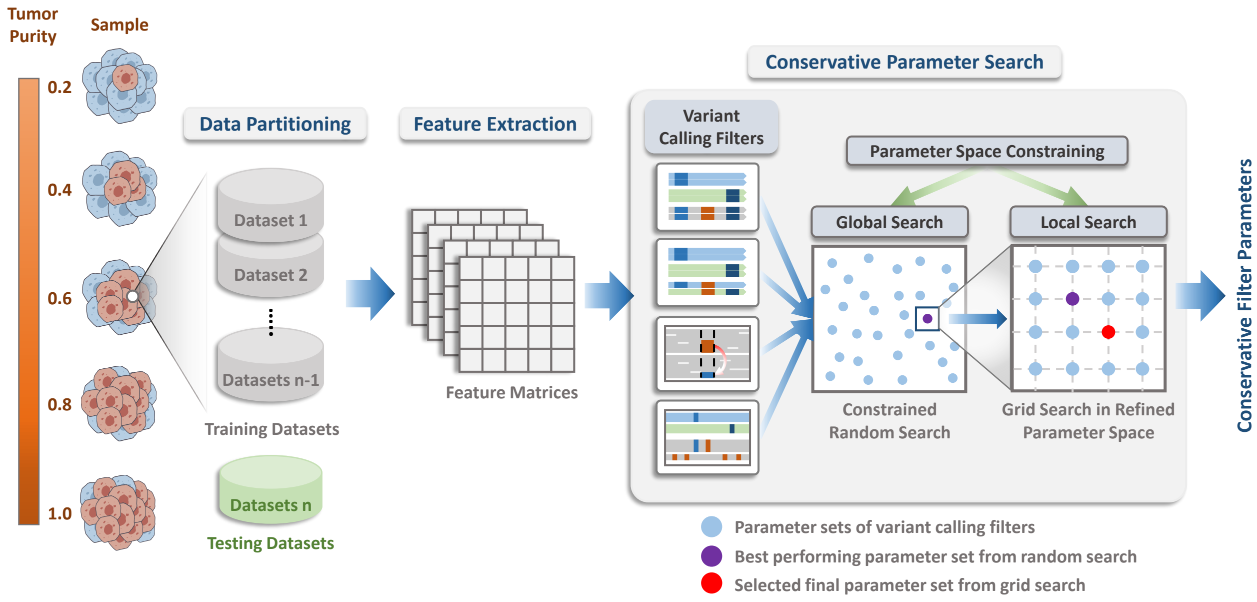

**Supplementary Figure 25**: Workflow of conservative parameter search across tumor purity levels, showing the data preparation steps and two-phase search strategy.

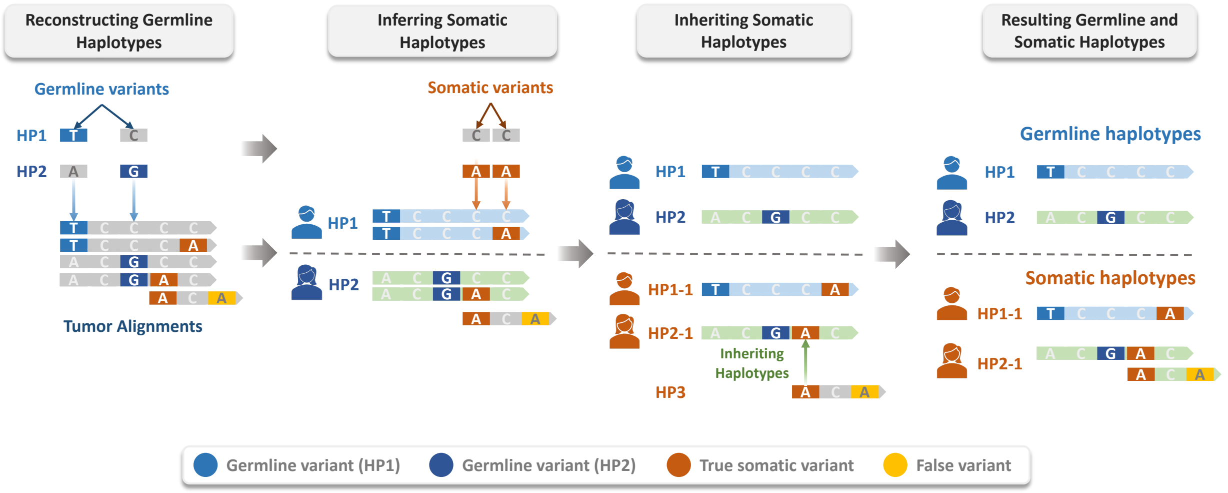

**Supplementary Figure 26**: Conceptual illustration of final somatic haplotagging, consisting of preliminary somatic haplotagging for initial variant classification, germline haplotype analysis for variant origin determination, and inheriting haplotype mechanism for processing reads lacking germline alleles (HP3 reads).

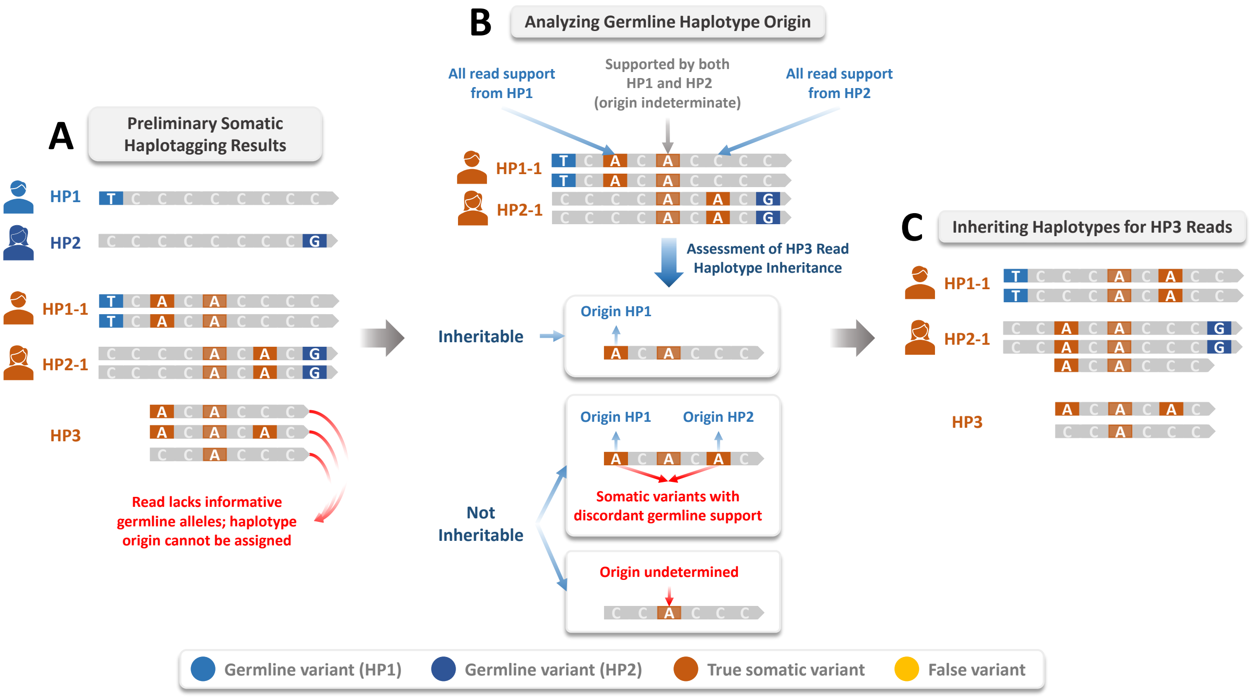

**Supplementary Figure 27**: Illustration of the inheriting haplotype mechanism for reads lacking germline alleles. (A) Preliminary somatic haplotagging results showing the initial classification of germline variants (HP1, HP2) and somatic variants (HP1-1, HP2-1, HP3), where HP3 reads lack informative germline alleles. (B) Germline haplotype origin analysis, examining whether somatic variants receive read support exclusively from HP1, exclusively from HP2, or from both haplotypes. (C) Final haplotype assignment results after applying the inheriting mechanism, where HP3 reads inherit the germline origin when consistent single-haplotype support is observed, or remain unassigned when support is discordant or indeterminate.

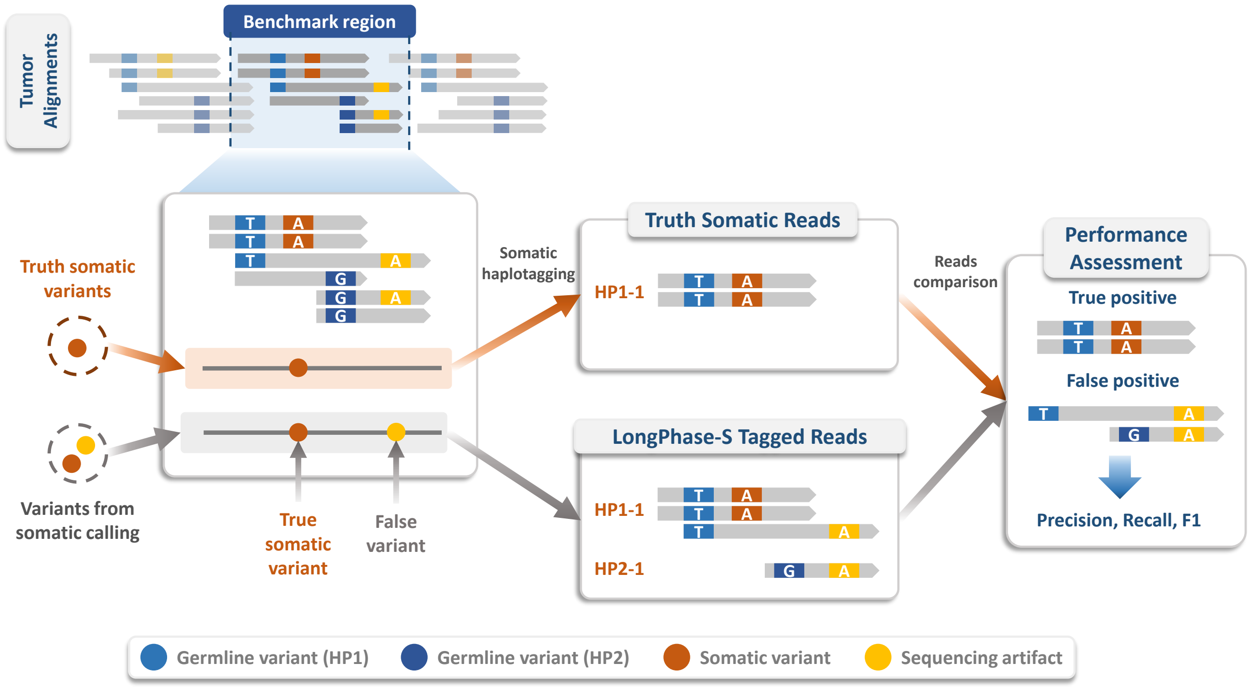

**Supplementary Figure 28**: Schematic of the read-level benchmarking framework for somatic haplotagging. This diagram outlines how LongPhase-S performance is evaluated by comparing somatic reads it tags against those derived from benchmark truth sets, enabling the calculation of precision, recall, and F1-score at the individual read level.

### 3 Supplementary Tables

**Supplementary Table 1**: Cell line datasets and corresponding benchmark truth sets from multiple sequencing sources (HKU, NYGC, ONT, UCSC, and DeepSomatic), with normal and tumor coverages.

| Dataset | Cancer Type | Source | Benchmark | Normal | Tumor |
|---|---|---|---|---|---|
| HCC1395_HKU | Breast ductal carcinoma | HKU | SEQC2 | 45.70x | 76.27x |
| HCC1395_NYGC | Breast ductal carcinoma | NYGC | SEQC2 | 29.14x | 79.52x |
| COLO829_ONT | Malignant melanoma | ONT | NYGC | 39.83x | 33.45x |
| COLO829_NYGC | Malignant melanoma | NYGC | NYGC | 45.85x | 108.49x |
| HCC1954_UCSC | Breast ductal carcinoma | UCSC | DeepSomatic | 34.03x | 82.44x |
| HCC1937_UCSC | Breast ductal carcinoma | UCSC | DeepSomatic | 25.68x | 158.44x |
| H2009_UCSC | Non-small-cell lung cancer | UCSC | DeepSomatic | 30.07x | 106.92x |
| H1437_UCSC | Non-small-cell lung cancer | UCSC | DeepSomatic | 49.28x | 79.10x |

**Supplementary Table 2**: Cross-validation performance metrics of MAE and $R^2$ (mean ± standard deviation) across training and testing sets.

| K | MAE (mean ± std) | | $R^2$ (mean ± std) | |
|---|---|---|---|---|
| | Training | Testing | Training | Testing |
| 1 | $0.0295 \pm 0.0022$ | $0.0401 \pm 0.0178$ | $0.9796 \pm 0.0032$ | $0.9603 \pm 0.0333$ |
| 2 | $0.0288 \pm 0.0035$ | $0.0404 \pm 0.0130$ | $0.9803 \pm 0.0049$ | $0.9591 \pm 0.0261$ |

**Supplementary Table 3**: Software and version used in the study.

| Category | Tool | Version |
|---|---|---|
| Somatic Variant Calling | ClairS (SS+RS model) | v0.4.1 |
| | ClairS (SS model) | v0.4.1 |
| | DeepSomatic | v1.8.0 |
| Germline Variant Calling | Clair3 | v1.0.10 |
| Read Alignment | Minimap2 | 2.24-r1122 |
| BAM Subsampling | Samtools | v1.13 |
| Coverage Calculation | Mosdepth | v0.3.3 |
| Variant Call Benchmarking | som.py | v0.3.12 |
